# T6SS-associated Rhs toxin containers: Structural and functional insights into bacterial weaponry and self-protection

**DOI:** 10.1101/2024.02.26.582050

**Authors:** Claudia S. Kielkopf, Mikhail M. Shneider, Petr G. Leiman, Nicholas M.I. Taylor

## Abstract

Bacteria use the type VI secretion system (T6SS) to secrete a variety of toxins into pro- and eukaryotic cells via a machinery consisting of a contractile sheath and a rigid tube. Rearrangement hotspot (Rhs) proteins represent one of the most common T6SS-secreted effectors. The C-terminal toxin domain of Rhs proteins displays great functional diversity, while the large Rhs core is characterised by YD repeats. T6SS- associated Rhs proteins are attached to the VgrG spike protein, often via an N- terminal PAAR-repeat domain. Using X-ray crystallography, we here elucidate the Rhs core structures of PAAR- and VgrG-linked Rhs proteins from *Salmonella bongori* and *Advenella mimigardefordensis*, respectively. The Rhs core forms a large container made up of β-sheets that has a negatively charged interior and encloses a large volume. The presence of the toxin domain of *S. bongori* PAAR-linked Rhs does not lead to ordered density in the Rhs container, suggesting the toxin domain is at least partially unfolded. Together with bioinformatics analysis showing that Rhs toxins predominantly act intracellularly, this suggests that the Rhs core domain functions two-fold, as safety feature for the producer cell and as delivery mechanism for the toxin domain. Our results strengthen our knowledge of Rhs structure and function.

## Introduction

To gain an advantage over their competitors, bacteria employ various survival strategies. One of these strategies is the type VI secretion system (T6SS) of Gram-negative bacteria. Ancestrally related to contractile tails of bacteriophages (Leiman et al. 2009), T6SS can inject effector proteins, e.g. hydrolase enzymes, across the cell membrane of eukaryotic or, more commonly, prokaryotic target cells (Green and Mecsas 2016, Pukatzki et al. 2006, Durand et al. 2014, Hood et al. 2010). The T6SS consists of a membrane-anchored contractile sheath that surrounds a tube (Leiman et al. 2009, Bonemann et al. 2009, Basler et al. 2012). The tube carries a trimeric spike protein, VgrG, which is “sharpened” by a pointy PAAR-repeat protein at its apex (Chang et al. 2017, Shneider et al. 2013). The contraction of the tail sheath structure results in the expulsion of the tail tube for protein secretion across the target cell membrane (Basler et al. 2012).

Transferred proteins typically have a toxic effect on target cells which can be eukaryotic cells (Hachani, Wood, and Filloux 2016, Pukatzki et al. 2006) but more often are bacteria of other or of sibling species (Hood et al. 2010, Coulthurst 2019). To prevent self-intoxication, the toxin-producing cell will therefore normally also express an immunity protein (Ruhe, Low, and Hayes 2020). These so-called toxin-antitoxin systems are immensely variable and play an important role in bacterial competition (Jurenas et al. 2022).

Unlike in other secretion systems, the secretion signal in T6SS had long been elusive and does not appear to reside in a short signal peptide sequence. Rather, proteins are secreted because they are associated either covalently or non-covalently with the tail tube-spike assembly (Alcoforado Diniz, Liu, and Coulthurst 2015, Shneider et al. 2013). The secretion selection signal is thus encoded in the three-dimensional structure of the effector.

For secretion, T6SS effectors can be either packaged into the lumen of the Hcp tube or attached to the VgrG-PAAR spike complex, similar to a poisoned arrowhead. In the latter case, the toxin domain forms a C-terminal extension of either VgrG or PAAR domain or an interaction with VgrG or PAAR. Spike tip interaction of effectors can be mediated by chaperones (Liang et al. 2015, Burkinshaw et al. 2018) and even by house-keeping proteins, such as the translation elongation factor Tu (Whitney et al. 2015).

One of the most common extensions of T6SS-associated PAAR domains are Rearrangement hotspot (Rhs) elements (PFAM03527) (Shneider et al. 2013). A typical Rhs genetic element contains at least four open reading frames (ORF): The first ORF encodes a > 1,000 amino acid protein with multiple YD (tyrosine-aspartate) repeats, followed by a small C-terminal toxin domain (about 100 residues). The second downstream ORF encodes a small immunity protein (Jackson et al. 2009). Further downstream, several pairs of toxin domain-immunity ORFs may be present, where the toxin ORF carrying a small fragment homologous to the C-terminal part of the YD-repeat sequence. Homologous recombination can create a new YD-repeat domain-toxin fusion, giving the Rhs family its name (Jackson et al. 2009, De Maayer et al. 2011).

Recent work on T6SS-associated Rhs proteins shows that the YD-repeat domain (termed Rhs core from here on) forms a large container-like structure composed of β-hairpins (Jurenas et al. 2021, Gunther et al. 2022, Tang et al. 2022). T6SS- associated Rhs proteins undergo autocleavage before and after the Rhs core domain at the conserved motifs PVxxxxGE and DPxGL, respectively. The N- and C- terminal cleavage products remain associated with the Rhs core (Pei et al. 2020, Jurenas et al. 2021, Gunther et al. 2022, Tang et al. 2022). Cleavage is essential for toxicity of Rhs in T6SS in *Aeromonas dhakensis*, but not for secretion (Pei et al. 2020).

Notably, the BC component of Tc and ABC toxins is homologous to T6SS- associated Rhs (Busby et al. 2013, Gatsogiannis et al. 2018). The Tc/ABC-associated Rhs also undergoes cleavage after the DPxGL motif, and the core- encapsulated toxin domain is released through a change in pH (Busby et al. 2013, Meusch et al. 2014).

In addition to being PAAR-linked or PAAR-adaptor-bound, Rhs proteins are also known to associate with VgrG via direct or indirect interaction (Pei et al. 2020). However, no VgrG-linked Rhs proteins have been experimentally described to date. Furthermore, Rhs proteins remain functionally poorly characterised, despite their ubiquity. While Rhs proteins are important for toxicity and function as a quality control for T6SS assembly (Alcoforado Diniz and Coulthurst 2015, Donato et al. 2020), it remains unclear whether Rhs proteins are secreted in full and how toxin release takes place.

Here, we report the existence of genetically VgrG-linked Rhs and show that proteolysis occurs between VgrG and Rhs. To gain more insight into the structure and function of Rhs proteins, we determined the structure of the Rhs domains of both PAAR- and VgrG-linked Rhs proteins using X-ray crystallography. In agreement with previous findings, we show that the C-terminal toxin domain is cleaved but appears to be trapped inside the Rhs core. Using bioinformatics, we find that Rhs effectors act predominantly in the cytoplasm whereas other T6SS effectors act both in the cytoplasm and in the periplasm of target cells. Lastly, we investigate the occurrence of transmembrane domains (TMDs) and the VgrG2-interacting region (VIR) in Rhs effector sequences to probe the translocation mechanism of the effector into the target cell cytoplasm. Our analysis finds no consistent presence of TMDs or VIRs in Rhs effectors, pointing towards differences in translocation mechanisms.

## Results

### Bioinformatics analysis shows that T6SS-associated Rhs proteins are genetically fused to PAAR and VgrG domains

To gain insight into the Rhs protein family, we investigated it using bioinformatical methods. Searching InterPro for domain architectures, it was readily apparent that a very large number of Rhs proteins (2,337 proteins) carry a PAAR-repeat domain at their N-terminus, as has been described earlier (Shneider et al. 2013). However, we found a small number of Rhs proteins (107 Rhs proteins) that encode a complete VgrG protein at their N-terminus. This was unexpected as all known VgrG trimers bind one monomer of PAAR-linked Rhs, which results in a single copy of Rhs core in the spike complex. The existence of VgrG-Rhs fusions suggests that these proteins form a spike containing three Rhs molecules per spike. To structurally understand the similarities and differences of PAAR- and VgrG-linked Rhs cores, we determined the structure of representatives of both.

### PAAR-linked Rhs core domain folds into a container-like structure

We solved the structure of the PAAR-linked Rhs domain of the *S. bongori* S5MXP0 protein (ORF A464_2174), residues 360-1420 with an N-terminal GSGS extension (Figure 1 A and Figure S1 A – B). The protein crystallised in the P2_1_ space group with two molecules in the asymmetric unit (Table S1) and its structure could be solved by single isomorphous replacement with anomalous scattering using a Hg^2+^ derivative.

**Figure 1:**
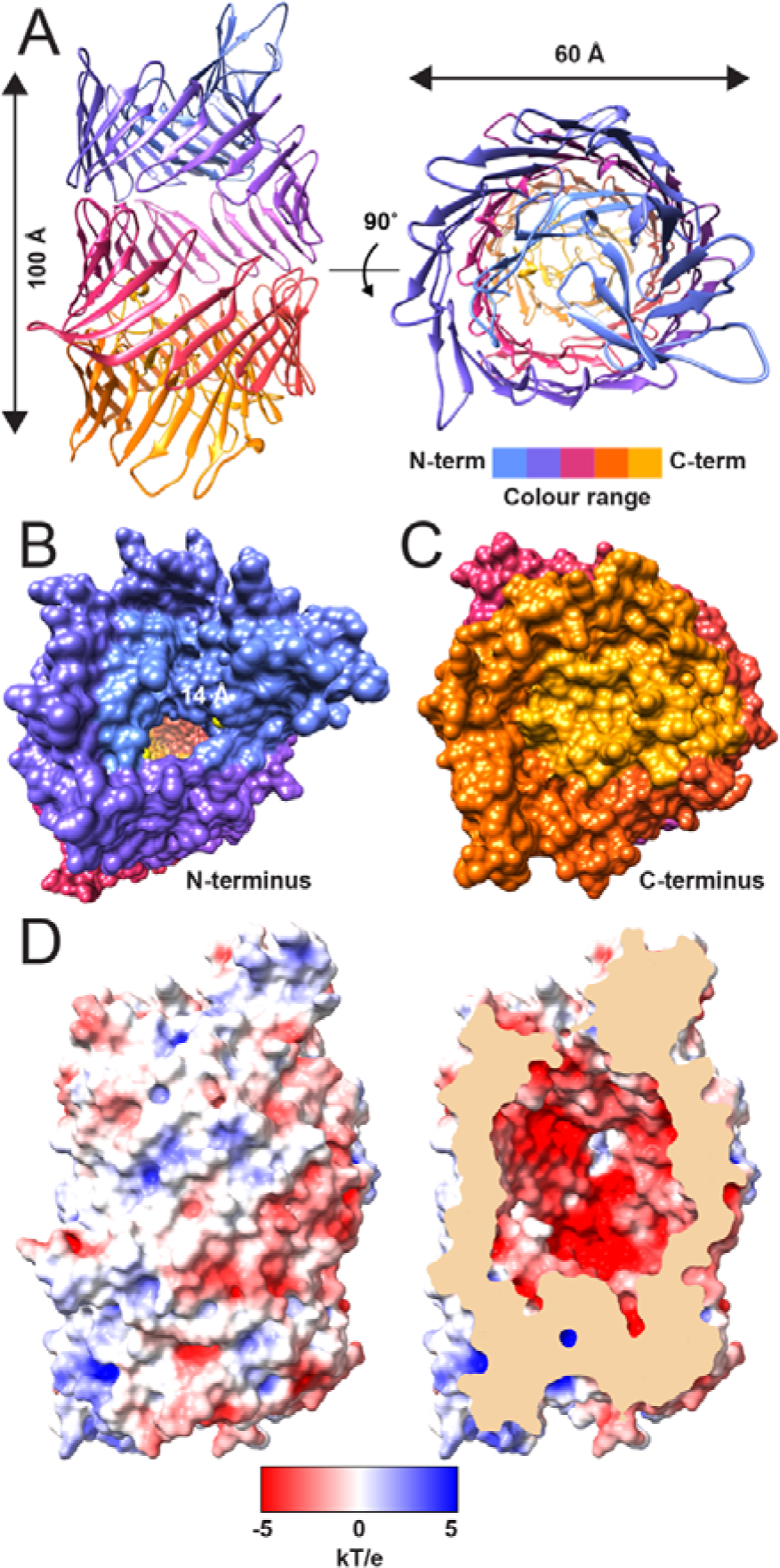
PAAR-linked Rhs core structure from *S. bongori*. A) Rhs core structure model in a side and top view, coloured from N-terminus (blue) to C-terminus (yellow). The Rhs core domain creates a container of 100 Å × 60 Å, creating an inner volume of ∼ 25,000 Å^3^. B) At the N-terminus, an opening of ∼14 Å creates a crevice, whereas C) at the C-terminal end, the container is fully closed. D) Surface representation of Rhs core structure, coloured according to electrostatic potential, as calculated using APBS, in a side view and clipped to expose the negative container inside.

The Rhs core forms a large container-like structure, consisting almost exclusively of staggered, oppositely oriented β-hairpins, in which the conserved YD motif forms the transition from a β-strand to a sharp turn. The β-hairpins create a continuous, gently curving β-sheet, while the N- and C-termini of the Rhs core form tight spirals that seal the core from both ends. The C-terminal end of the Rhs core, to which the toxin domain is linked in full length Rhs, is located well inside the container. The dimensions of the Rhs core are around 100 Å × 60 Å on the outside, creating an internal volume of ∼25,000 Å^3^ as determined using CASTp (Tian et al. 2018) with a probe radius of 2.5 Å.

The N-terminal part of the core has a small opening, with a maximal atom-to-atom distance of around 14 Å (Figure 1 B). In contrast, the C-terminal part of the core appears completely closed (Figure 1 C). The outside of the core is in many areas negatively charged (Figure 1 D). Inside of the core, the surface is even more uniformly negatively charged over the whole internal surface (Figure 1 D).

### T6SS-associated Rhs core structures from various species closely resemble each other

Three Rhs core structures have been published recently, PAAR-linked Rhs from *Photorhabdus laumondii* (Jurenas et al. 2021), PAAR-linked Rhs from *Pseudomonas protegens* (Gunther et al. 2022) and RhsP from *Vibrio parahaemolyticus* (Tang et al. 2022). Comparing these sequences for the Rhs core to the *S. bongori* Rhs core domain, the sequence identities range from 21 to 29%, and the sequence similarities range from 35% to 42%. Despite low sequence similarities, these structures closely resemble the *S. bongori* Rhs structure presented here with small differences in their loop regions. Loops formed by residues 796-802, 935-945, and 1120-1199 of *S. bongori* Rhs are larger than those in other Rhs cores. In other areas, most notably in the loop formed by residues 1312-1315 in *S. bongori*, the loops of *Pseudomonas* and *Photorhabdus* Rhs proteins are larger, with *Vibrio* RhsP being the same length as in *S. bongori* Rhs.

In addition, the N- and C-terminal regions of all four Rhs cores, which plug both ends of the core and might be important for toxin release, are structurally very similar. Interestingly, similar to the negative charge of the outside and inside surfaces of the PAAR-linked Rhs core of *S. bongori*, the *Pseudomonas* and *Photorhabdus* PAAR- linked Rhs as well as the uncleaved form of *Vibrio* RhsP are overall negatively charged (Figure 1 D, Figure S2 A – C). Only the cleaved form of *Vibrio* RhsP has a positively charged patch close to the hypothesised exit crevice (Figure S2 D, indicated with an arrow).

The structure of *S. bongori* T6SS-associated Rhs presented here is also structurally similar to that of the BC component of the ABC-toxin family (Busby et al. 2013, Meusch et al. 2014, Gatsogiannis et al. 2018). Residues 360 to 745 of *S. bongori* Rhs cover the B component part of the core, albeit not the β-propeller part of the B component that interacts with the A component and allows for toxin release through a conformation change (Meusch et al. 2014, Gatsogiannis et al. 2018). Rhs residues 746 to 1420 cover the C^NTR^ component. Unlike T6SS-associated Rhs proteins, the BC component consists of two proteins, although the BC proteins occur as apparent fusion protein in *Burkholderia rhizoxinica* (Busby et al. 2013). The internal volume of the Rhs core domain is smaller than the volume of the BC component from *Yersinia entomophaga* (Busby et al. 2013) (∼25,000 Å^3^ vs ∼37,000 Å^3^, respectively). In contrast to the BC component, the outside surface of T6SS-associated Rhs proteins is very smooth and lacks protrusions and structural “decorations”.

### Extended N-terminus in PAAR-linked Rhs core does not close the container

To test whether an extension to our original construct of *S. bongori* Rhs (residues 360-1420) would close the N-terminal opening of the core, we created and solved the crystal structure of a larger construct (residues 348-1420). However, the additional residues did not close the opening and instead appeared to be located inside the core, near residue 360 (Figure S3 A). The fragment is mostly disordered, which manifests in a weak and uninterpretable density (Figure S3 B). Comparing to an AlphaFold2 structure prediction, we can see that the N-terminal end of the plug region is indeed predicted to be located where the crevice is in our experimental structure, effectively closing off the Rhs container (Figure S3 C). A comparison with the structure of *Pseudomonas protegens* PAAR-Rhs (Gunther et al. 2022) shows that its “anchor” helix corresponds to an unstructured loop in our *S. bongori* PAAR- linked Rhs. Notably, PAAR-linked Rhs of *Photorhabdus laumondii* (Jurenas et al. 2021) also forms a loop in this region instead of a helix (Figure S3 D). This area was not modelled in RhsP from *Vibrio parahaemolyticus* (Tang et al. 2022).

### VgrG-linked Rhs core shows small differences to PAAR-linked Rhs

Recombinantly expressed VgrG-Rhs of *A. mimigardefordensis* (DPN7T strain, AHG65567 protein residues 1-1875) is cleaved into two parts, which correspond to the VgrG spike and Rhs core (Figure S1 F). The two parts form a complex that is stable enough to withstand a metal-affinity resin, TEV cleavage, anion exchange chromatography and size exclusion with several dialysis steps (Figure S1 D – F). In size exclusion chromatography, the complex has an apparent molecular weight (MW) of 691 kDa. This matches a combined MW of three full-length chains (228 kDa × 3 = 684 kDa), suggesting that the complex is composed of a trimeric VgrG spike and three Rhs cores (Figure S1 D and E).

We pursued crystallisation trials of the VgrG-Rhs core complex and obtained crystals that diffracted to 3.4 Å resolution. However, after solving the structure by molecular replacement using the *S. bongori* PAAR-Rhs core as a search model, it became apparent that no VgrG was present in these crystals. Instead, the Rhs domain crystallised in the P1 space group with four molecules in the asymmetric unit (Table S1).

The overall structure of the VgrG-linked Rhs core is very similar to that of the PAAR- linked Rhs, with some small differences (Figure 2 A). First, residues 1141-1204 of PAAR-linked Rhs form four β-strands, while the corresponding region of VgrG-linked Rhs (residues 829-847) forms mostly a big loop with only one short β-strand (Figure 2 B). Second, several loops are longer in PAAR-linked Rhs (Figure 2 A and B). Third, the VgrG-linked Rhs core is fully closed by its N-terminal region that occupies the open crevice of PAAR-linked Rhs (Figure 2 A, black arrow, and C). Like PAAR- linked Rhs, VgrG-linked Rhs is highly negatively charged, both on the outside and on the inside (Figure 2 D). Like RhsP and PAAR-linked Rhs of *S. bongori* and *Photorhabdus laumondii*, VgrG-linked Rhs is lacking an anchor helix (Figure S3 D).

**Figure 2:**
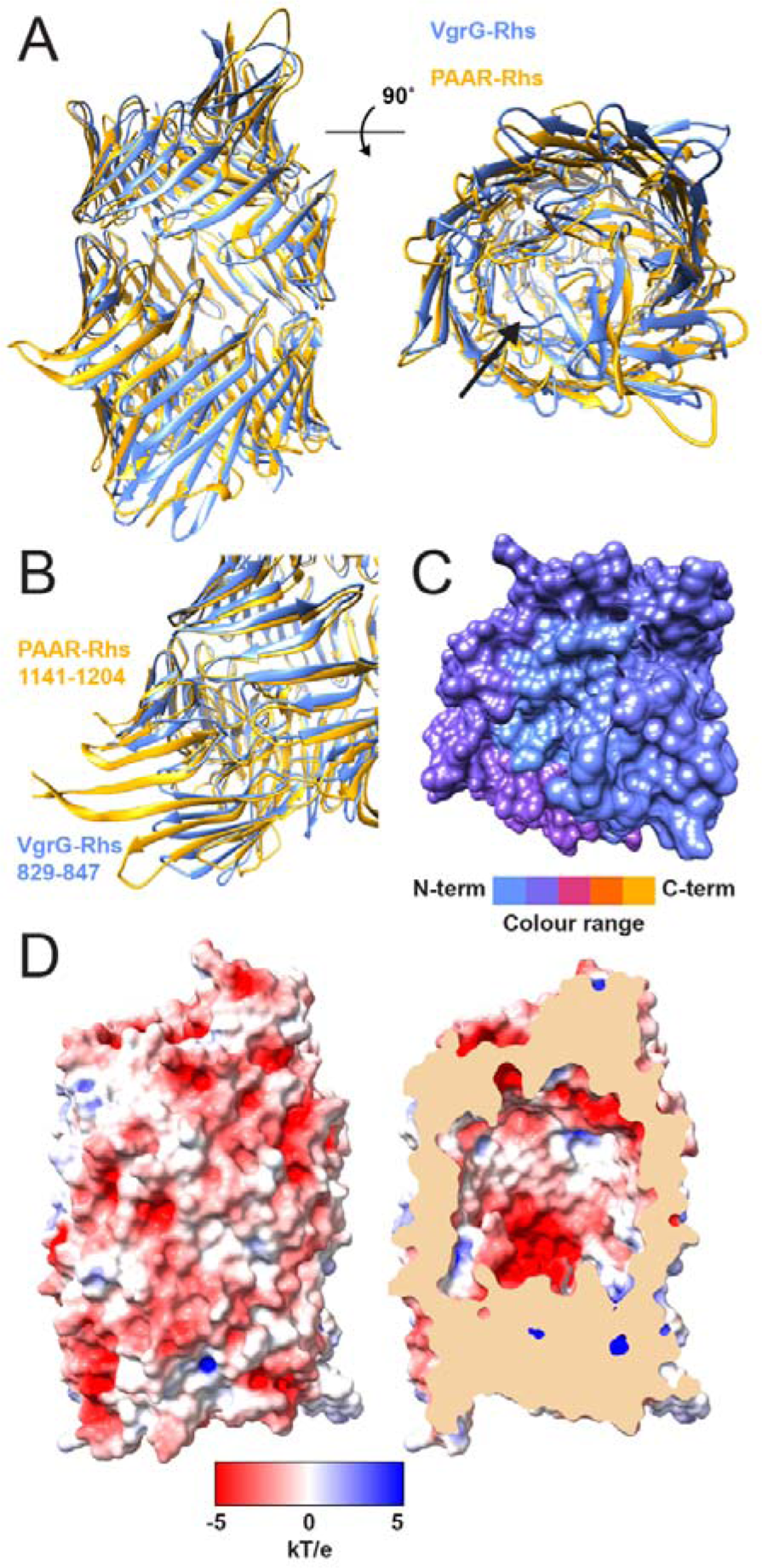
VgrG-linked Rhs structure from *A. mimigardefordensis*. A) VgrG-linked Rhs core structure (blue) in the side and top view, superimposed onto PAAR-linked Rhs (yellow). The two structures are virtually identical. No VgrG molecule was present in the *A. mimigardefordensis* crystal. B) Several loops are more extended in the PAAR-linked Rhs compared to VgrG-linked Rhs. C) VgrG-linked Rhs appears completely closed at the N- terminus, in contrast to PAAR-Rhs from *S. bongori*. D) Surface representation of VgrG-linked Rhs, coloured according to electrostatic potential, as calculated using APBS, showing a highly negatively charged surface, both on the outside of the container as well as on the inside, similar to PAAR-Rhs.

To verify cleavage of the *A. mimigardefordensis* Rhs protein at the C-terminus before the toxin, we also cloned the full-length AHG65567 gene and expressed and purified the protein. On SDS-PAGE, a clear band is detectable at ∼16 kDa (Figure S1 G), which corresponds well with the cleaved C-terminal toxin domain (residues 1876- 2012, 15.9 kDa MW).

As the resolution of the VgrG-linked Rhs dataset was lower than for the PAAR-linked Rhs (Table S1, Figure S4 A – D), we compared the crystal structure to an AlphaFold2-prediction (Figure S4 E and F). The AlphaFold2 prediction agreed with the crystal structure remarkably well, including the conformation of potentially flexible loops with a single exception. The structure of residues 800-829 is different from that of the AlphaFold2 prediction (Figure S4 F). The crystal structure shows a β-strand, while AlphaFold2 predicts a big loop.

### Rhs core-encapsulated toxin is likely in a molten globule state or unfolded

The most interesting open question regarding Rhs proteins is arguably the mechanism of toxin release. To gain insight into this question, we produced recombinant PAAR-linked Rhs with its cognate toxin (residues 360-1530). This required co-expression of its respective immunity protein (uniprot S5MXF1, ORF A464_2173). On SDS-PAGE, the purified Rhs core-toxin complex shows a clear band for the toxin in addition to the Rhs core band, indicating autocleavage of the toxin domain (Figure S1 C). The complex was crystallised in conditions that were nearly identical to those of the empty Rhs core, indicating that the toxin is fully buried inside the Rhs core. The structure of the Rhs-toxin complex was then solved by MR. However, the Rhs core contained no additional ordered density inside the core (Figure 3 A − C). Checking crystal packing, we verified that there is no unassigned density outside of the Rhs core that could correspond to the toxin domain (Figure S5 A − F).

**Figure 3:**
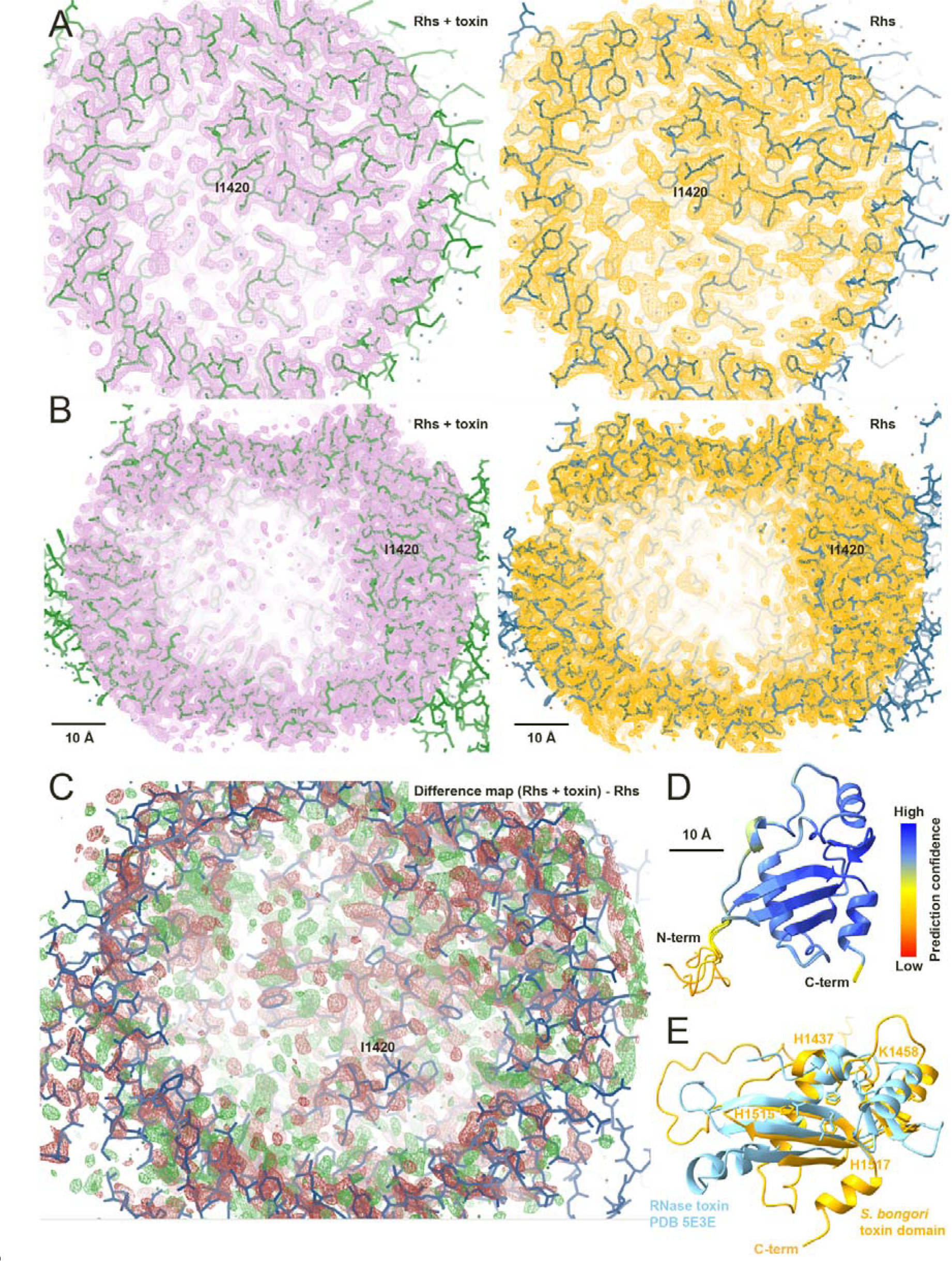
PAAR-linked Rhs with cognate toxin does not show additional density inside the Rhs container. A) The structure solved for PAAR-Rhs + toxin (left) from *S. bongori* does not show any extra density inside the container that could correspond to the toxin, in comparison to PAAR-Rhs alone (right). The last traceable residue, I1420, is indicated and the image is viewing the inside of the container along the long axis towards the C-terminal end. B) Views of the inside of the Rhs container for PAAR-Rhs + toxin (left) and PAAR-Rhs (right) along the short axis show no additional density in PAAR-Rhs + toxin inside the container (scale bar for 10 Å shown). C) Difference map created by subtracting the Rhs map without toxin from the Rhs + toxin map. No additional density in Rhs + toxin (green) is present inside the container. The last traceable residue, I1420, is indicated and the image is viewing the inside of the container along the long axis towards the C-terminal end. D) AlphaFold2 predictions of the *S. bongori* PAAR-Rhs toxin domain show a well-folded structure with three α-helices and β-strands. Size-wise, the predicted structure fits into the container. Images B and D are shown at the same scale (scale bar for 10 Å shown). E) Overlay of *S. bongori* Rhs toxin domain (yellow) with RNase toxin CdiA-CT (blue), showing that the *S. bongori* Rhs toxin domain has an RNase fold. The experimentally determined active residues in CdiA-CT (H175, R186, T276 and Y278) and the equivalent residues in the *S. bongori* Rhs toxin domain (H1437, K1458, H1515 and H1517, labelled in the figure) are shown as stick representation. The contour level in A and B is 0.8 r.m.s.d.

### The toxin domain of S.bongori PAAR-Rhs is likely an RNase

The putative toxin of *S. bongori* PAAR-linked Rhs is 110 amino acids long. No protein family association could be found for the toxin part using the Conserved Domain Database (CDD), pfam and InterPro. Using HHpred with default parameters, only one significant homology could be detected to an RNA-binding protein VP-40 from Ebola virus. According to JPRED_PSSM, the toxin consists of 10% α-helical and 16.4% β-strand structure.

AlphaFold2 predicts the toxin to contain three α-helices and three long and two shorter β-strands (Figure 3 D). A short N-terminal stretch of the toxin shows low prediction confidence and is probably disordered. Importantly, the toxin is small enough to fit inside the Rhs core in its predicted folded state (∼ 35 Å x 35 Å, Figure 3 B and D are on the same scale). However, the absence of any ordered electron density corresponding to the toxin inside the core suggests that the toxin is in a molten globule state.

A DALI search of the toxin structure prediction against the PDB resulted in significant hits with nucleases and RNase toxins (Table S2). The top match was CdiA-CT (PDB 5E3E, Z-score 5.2, root mean square deviation 3.0 Å, 77 equivalent residues), an RNase toxin of a contact-dependent growth inhibition toxin from *Yersinia kristensenii* (Batot et al. 2017). Manual examination of the *S. bongori* Rhs toxin domain shows that it has an RNase fold (Figure 3 E and Figure S6 A and B). When comparing the toxin domain to the experimentally determined active sites of CdiA-CT (H175, R186, T276 and Y278), the equivalent residues in the *S. bongori* Rhs toxin are H1437, K1458, H1515 and H1517 (Figure 3 E).

### Most Rhs effectors are predicted to act in the cytoplasm

All T6SS-associated Rhs proteins are thought to carry an effector after the container part of the protein (Ma et al. 2017, Liu et al. 2020). We therefore performed bioinformatical analyses, aiming to decipher common characteristics of T6SS- associated Rhs effectors. We compiled predicted and confirmed T6SS effectors from the SecReT6 T6SS database, UniProt (PAAR-Rhs and VgrG-Rhs), and the list of effectors reported in Liu et al. (2020). In total, we identified 18,330 unique T6SS- associated effector proteins (Table S3 and supplementary FASTA file). We then determined conserved domains in these effectors using CDD search to identify prefix domains (PAAR, VgrG, Rhs, Hcp, and VipA as a negative control) and effector functions (Figure 4 A, Tables S3 − S5). We found that 5,541 effectors contain an Rhs domain, and thereof, 4,743 are PAAR-linked and 107 are VgrG-linked Rhs proteins (Figure 4 A).

**Figure 4:**
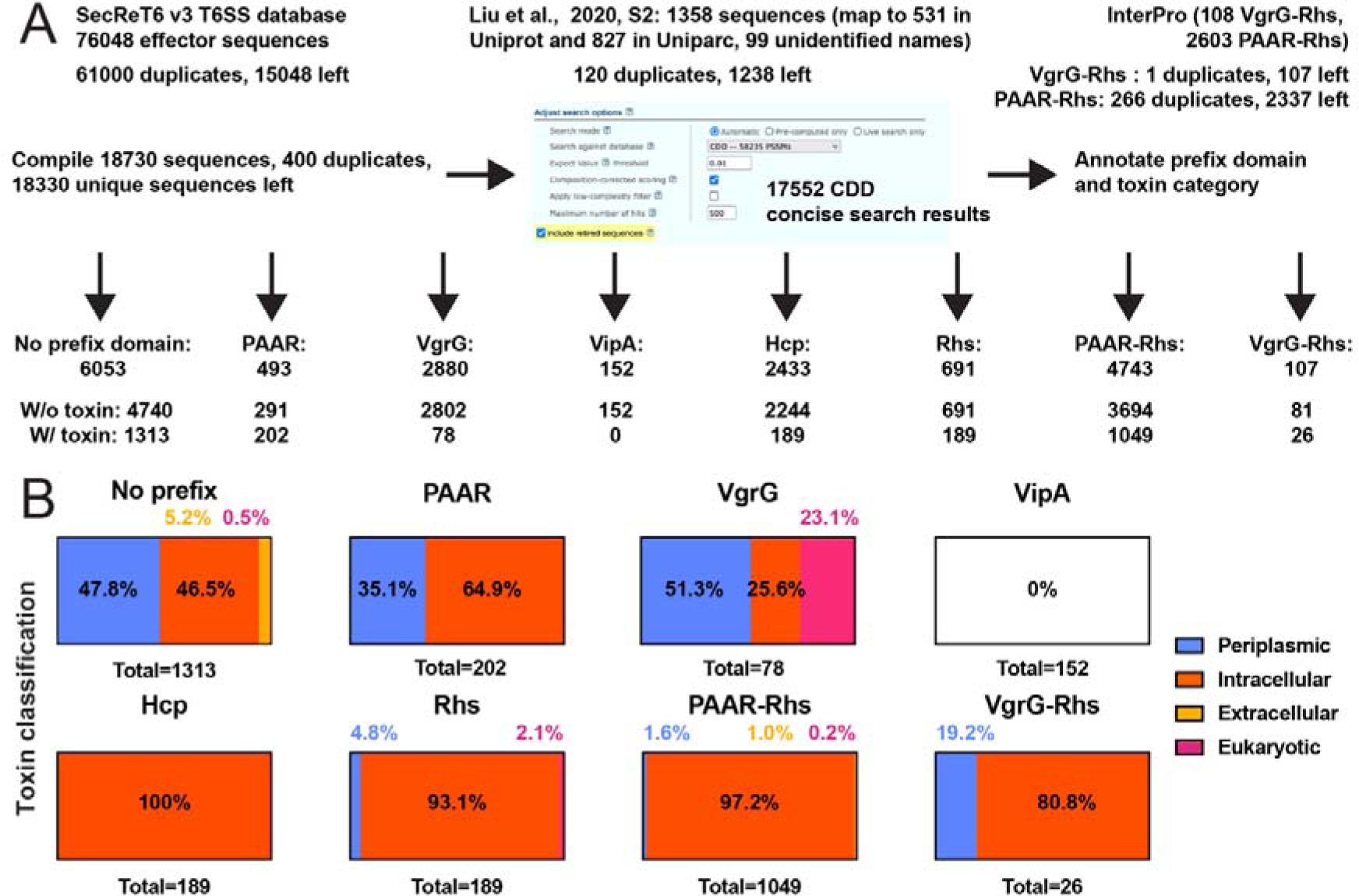
T6SS effector analysis. A) Workflow for effector analysis. T6SS effector sequences are collected, duplicates by sequence removed and annotated using CDD. B) Analysis of annotated effector domains per prefix domain. Rhs proteins act predominantly intracellularly.

We next investigated differences in MWs and isoelectric points of Rhs and non-Rhs effectors (Figure S7) for full-length proteins, as well as for their effector domains by retaining the sequence after the conserved cleavage motif or their annotated toxin domain, respectively. Full-length Rhs-containing proteins are on average larger than other effector-carrying proteins (Figure S7 A), due to their massive YD-repeat domain. VgrG-linked effectors also have high MWs. However, there are 43 Rhs proteins with a MW less than 100 kDa. They contain between 2 and 16 YD repeats before the C-terminal toxin domain. AlphaFold2 predictions of these short Rhs proteins show that they contain a full-length effector domain, but only a part of the Rhs core domain (as few as four β-strands) that cannot envelope the effector (Figure S8 A and B). For some of these, a complete Rhs core, including a toxin domain, is encoded by a nearby gene (e.g. UniProt entry X2H4X5 for UPI00042E4AAB in our toxin list). It is likely that these effectors are part of the of diversification mechanism of effectors through recombination and not expressed (Jackson et al. 2009, De Maayer et al. 2011). Analysis of isoelectric points of complete effector sequences shows that Rhs effectors mostly have an acidic pI and so do VgrG-linked effectors (Figure S7 B). The isoelectric points of effectors without a prefix domain, PAAR- linked effectors without an Rhs domain and Hcp effectors are bimodally distributed, with acidic and basic isoelectric points. In general, protein isoelectric points tend to be bimodally distributed, with bacterial proteomes being slightly more acidic (Kiraga et al. 2007).

Rhs proteins are known to undergo autoproteolysis that separates the Rhs core from the effector domain. We therefore cleaved all Rhs effectors in silico at the consensus DPxG[LWF] motif. For comparison, we extracted the annotated C-terminal effector domains of PAAR- and VgrG-linked effectors and investigated MW and isoelectric points (Figure S7 C and D). The MWs of Rhs-linked effectors without VgrG or PAAR domain cluster into two groups, one group with small MWs as PAAR- and VgrG-linked Rhs proteins exhibit, and another group of proteins with a larger MW of ∼38 kDa (Figure S7 C). PAAR-linked effectors without Rhs are on average about as small as the Rhs-linked effectors. VgrG-linked effectors tend to have larger MWs. One could speculate that Rhs-linked toxin domains have a limited size range since they need to fit inside the Rhs container, but there is the subgroup of Rhs-linked effectors without VgrG or PAAR with a larger MW of ∼38 kDa. Interestingly, this subgroup consists almost exclusively of toxin domains from the *Vibrio* genus. The isoelectric point distributions of all cut effector domains are bimodal, with acidic and basic isoelectric points (Figure S7 D), as would be expected proteome-wide (Kiraga et al. 2007).

T6SS effectors disrupt or dysregulate various cellular functions and thus, their sites of action and destinations vary greatly. We therefore categorised our list of T6SS effectors by assigning a functional mechanism and location to each effector function (Figure 4 B and Tables S3 and S5). We found that effectors without prefix domains are predicted to act both intracellularly and periplasmically, whereas PAAR effectors function more often intracellularly. About a half of VgrG effectors are predicted to act periplasmically, and about a quarter each are intracellular and eukaryotic. In contrast, Rhs and PAAR-linked Rhs effectors are predicted to operate almost exclusively intracellularly, for example as nucleases and deaminases (Figure 4 B and Table S3). VgrG-linked Rhs effectors are also predicted to act predominantly intracellularly, but around 20% are periplasmic. We also analysed the T6SS VipA sheath proteins for the presence of effector domains and found none, which supports the validity of the prediction approach. Furthermore, all Hcp-linked effectors have intracellular functions, as expected from T6SS tube proteins.

### TMDs and VIRs are not consistently present in Rhs effectors

One suggested mechanism for Rhs effector delivery into the target cell involves translocation across the inner membrane, with the help of an N-terminal transmembrane domain (TMD) (Jurenas et al. 2021, Gunther et al. 2022, Quentin et al. 2018). We investigated if Rhs effectors in our list have a predicted TMD, using an algorithm that others used in the past, namely TMHMM (Krogh et al. 2001) as well as its successor DeepTMHMM (Hallgren et al. 2022). We found that only a very small percentage of the Rhs and VgrG-linked Rhs proteins is predicted to contain a TMD (Table 1). While the two algorithms do not agree for PAAR-Rhs TMD predictions, there are PAAR-linked Rhs effectors for which neither algorithm predicts a TMD.

**Table 1:** Predicted TMDs in Rhs effectors.

|  | **Rhs (closed core,**<br>**138 proteins)** | **PAAR-Rhs**<br>**(1038 proteins)** | **VgrG-Rhs**<br>**(26 proteins)** |
| --- | --- | --- | --- |
| TMHMM | 6 (4.3%) | 702 (67.6%) | 1 (3.8%) |
| DeepTMHMM | 4 (2.9%) | 7 (0.7%) | 1 (3.8%) |
| Thereof concurring | 1 (0.7%) | 4 (0.4%) | 1 (3.8%) |
| Predicted TMDs<br>overlap | No | 2 no, 2 partially | Partially |

It has also been hypothesised that a specific region of *Vibrio parahaemolyticus* RhsP, which is termed VgrG2-interacting region (VIR) and is located right before the effector domain, plays a role in toxin release. A conformational change in VIR after autocleavage promotes Rhs core dimerisation and the escape of the toxin domain through an opening on the Rhs core side (Tang et al. 2022). To check for the presence of VIR sequences in other Rhs proteins, we created a hidden Markov model (HMM) of VIR using jackhmmer and the VIR from *Vibrio parahaemolyticus* (Supplementary files 6 and 7) and ran hmmsearch on all Rhs-containing sequences for which we could identify a toxin domain. We found 66 matches in other Rhs proteins from the *Vibrio* genus with sequence identities ranging from 100% to 79%, but not in Rhs of other species, in PAAR-Rhs, or VgrG-Rhs (Supplementary file 8). Interestingly, these matches are also the toxin domains that we found to have a larger MW (Figure S7 C). No non-Rhs sequence in our sequence compilation matched the HMM of VIR. In the jackhmmer result, three Rhs proteins of *Shewanella* and one Rhs protein of *Pseudoalteromonas* matched the VIR HMM (Supplementary files 6), showing that VIR is not exclusive to the *Vibrio* genus, but is not widespread among Rhs proteins.

## Discussion

Rhs proteins are among the most common effectors secreted by the T6SS (Koskiniemi et al. 2013). However, despite recent work aimed at the elucidation of their structure and mechanism of action (Pei et al. 2020, Jurenas et al. 2021, Gunther et al. 2022, Tang et al. 2022), key aspects of effector delivery and release remain unknown.

Here, we present crystal structures of two different Rhs protein cores, from *S. bongori* and *A. mimigardefordensis*, including a structure of Rhs core with a packaged effector domain. Both Rhs cores form a large container-like structure with a negatively charged cavity (Figures 1 and 2). Both structures are very similar to Rhs cores published recently (Jurenas et al. 2021, Gunther et al. 2022, Tang et al. 2022). Neither of the structures described here contains an anchor helix, which has been proposed to be responsible for keeping the N-terminal cleavage site and the plug domain in place at the Rhs core after autocleavage (Gunther et al. 2022). Our structures, as well as the structures by Jurenas et al. (2021) and Tang et al. (2022), show the plug domain in place without an anchor helix. Hence, the loop that covers this area appears to be sufficient to keep the plug domain in place (Figure S3 D).

Many Rhs proteins carry a PAAR domain (Shneider et al. 2013), as does the *S. bongori* Rhs described here. We show that Rhs proteins are also genetically fused to VgrG, for example the *A. mimigardefordensis* Rhs (Figure 4). Such a “megaspike” with three Rhs proteins per trimeric VgrG carries three effectors that are delivered to a target cell in a single T6SS firing event (schematic image in Figure S1 H). Although VgrG-effector fusions are not uncommon and such an effector was one of the first ever described in a T6SS (Pukatzki et al. 2007), such effectors target eukaryotic proteins (e.g. by crosslinking actin) or membrane/periplasm-associated components (e.g. the peptidoglycan) (Pukatzki et al. 2007, Carobbi et al. 2022).

The presence of the *S. bongori* PAAR-Rhs effector in the Rhs core did not give rise to any ordered density inside the Rhs core in our crystal structure (Figure 3). We could not find any unassigned density outside of the Rhs core, either (Figure S5). In the recently published PAAR-linked Rhs structures, the cognate toxins were also expressed, but no clear density could be assigned them, either (Jurenas et al. 2021, Gunther et al. 2022). The same is true for RhsP from *Vibrio parahaemolyticus*, where the linker domain between the Rhs core and the toxin is ordered but not the toxin itself (Tang et al. 2022). Together, this suggests that the effector domain remains partially or entirely unfolded inside the Rhs core. Given the size of the internal Rhs cavity and our MW analysis showing that Rhs toxin domains are as large as 38 kDa (Figure 1 and Figure S7), it is likely that the Rhs core can encapsulate larger, engineered payload.

Our analysis shows that Rhs effectors act predominantly intracellularly (Figure 4). Additionally, a body of evidence in the literature shows that intracellular T6SS effectors often contain Rhs repeats (Zhang et al. 2012, Lien and Lai 2017, Repizo et al. 2019). Taken together with the observation of an unfolded toxin domain inside the Rhs core, these results suggest two important functions of the Rhs core.

The first suggested function of the Rhs core is to protect the producer cell from the toxic effect of the effector domain. However, the core by itself is likely insufficient for the protection because we failed to obtain clones of full-length Rhs without the immunity protein. As our results show, the Rhs core is made entirely of β-hairpins that interact sequentially to form a shell-like structure. The formation of the secondary and tertiary structure can therefore take place simultaneously with the synthesis of the Rhs polypeptide chain. The Rhs core can thus co-translationally encapsulate the effector domain and shield it from the cytosol of the producer cell. Additionally, by keeping the effector in an unfolded or molten globule state, as our results and those from other suggest (Jurenas et al. 2022, Gunther et al. 2022, Tang et al. 2022), the Rhs core effectively protects the producer cell from the effector.

The second suggested function of the Rhs core is to contribute to delivering the effector into the target cell cytoplasm where is exerts its toxin function. However, the exact mode of translocation is elusive, and it is unknown if the whole Rhs core with effector or the effector only is translocated into the cytoplasm.

For Jurenas et al. (2021) and Gunther et al. (2022), the proposed toxin delivery and release mechanism for PAAR-linked Rhs hinges on the presence of a TMD, which is hypothesised to insert itself into the inner membrane of the target cell. Our results, however, show that TMHMM predicts a TMD in many PAAR-Rhs in our list, but not in all of them. Similarly, no TMD is predicted in other Rhs proteins (Table 1). TMHMM’s successor algorithm DeepTMHMM, which is as good as TMHMM at predicting α-helical TMDs and better at predicting signal peptides and β-barrel TMDs (Hallgren et al. 2022), predicts that hardly any Rhs protein contains a TMD (Table 1). Hence, there is either a difference in delivery mechanism depending on whether an Rhs protein has a TMD or not, or there is a different delivery mechanism that is independent of a predicted TMD.

Tang et al. (2022) propose an entirely different mechanism for toxin delivery and release for Rhs proteins with a VIR. Either before or after delivery of Rhs into the target cell, but after autocleavage, the VIR domain undergoes a conformational change inside the Rhs core. VIR and the effector then leave the Rhs core through an opening on the side, which leads to Rhs dimerisation. We could find a VIR in other *Vibrio* Rhs proteins, but not widely in Rhs of other species or in PAAR-Rhs or VgrG- Rhs (Supplementary files 6 – 8). This observation raises the question if the suggested mechanism of Rhs toxins via VIR is specific to the *Vibrio* genus. On the other hand, Tang and colleagues point out that dimer formation in cleaved Rhs was also observed by Gunther et al. (2022). Jurenas et al. (2021) did not observe such dimer formation, which could be due to using Rhs with mutated cleavage motif for cryo-EM, which prevented autocleavage of the toxin domain. In our experiments, we observe two and four molecules per asymmetric unit for *S. bongori* PAAR-Rhs (also with extended N-terminus) and *A. mimigardefordensis* VgrG-Rhs, respectively. Both constructs cover the Rhs core only, so no autocleavage has occurred here, but the construct could mimic the cleaved Rhs. However, for the *S. bongori* Rhs core with the toxin, only one molecule per asymmetric unit was found (Table S1 and Figure S5). It is unclear if these observations hint at a mechanistic role of Rhs dimerisation or if it is caused by vitrification or crystal packing effects.

For Rhs proteins without VIR, the next step of the effector leaving the Rhs core is also unknown. Jurenas et al. (2021) hypothesise that conditions in the target cell contribute to releasing the effector domain, or that a conformational change in the Rhs core is required for toxin release. On the other hand, Gunther et al. (2022) speculate that translocation-induced pulling and spontaneous refolding of the effector causes the effector release. It is also interesting to compare T6SS- associated Rhs effectors to the insecticidal ABC toxins for which a conformational change in the β-propeller of the B component is responsible for toxin release (Busby et al. 2013, Meusch et al. 2014, Gatsogiannis et al. 2018). The rest of the B and C component which are the “counterpart” to the T6SS-associated Rhs core, however, display no conformational change upon toxin release (Gatsogiannis et al. 2018). Hence, the T6SS-associated Rhs core is likely to have evolved a different toxin release mechanism. The keen resemblance of the Rhs core structure to a peel-away wrapper with pre-determined breaking points in between the layers of Rhs repeats could have implications for the process of toxin release (Figure 1 A).

One could further speculate if the strong negative charge the Rhs core contributes to the functions of the Rhs core (Figures 1– 2 and S2), i.e., keeping the producer cell safe from the toxin, or translocating and releasing the toxin domain. Further experiments specifically addressing these aspects of the structure and function will be necessary to elucidate the remarkable properties of Rhs proteins.

## Methods

*Cloning, expression and purification of* S. bongori *Rhs core domain,* S. bongori *Rhs core domain (extended N-terminus), and* A. mimigardefordensis *DPN7T VgrG-Rhs* The cloning, expression and purification procedure was the same for three constructs: Rhs core domain of *S. bongori* N268-08 protein (uniprot S5MXP0, ORF A464_2174) (residues 360-1420), Rhs core domain with an extended N-terminus of *S. bongori* N268-08 protein (uniprot S5MXP0, ORF A464_2174) (residues 348-1420) and *A. mimigardefordensis* DPN7T strain VgrG-Rhs protein (uniprot W0PIA9, AHG65567, ORF MIM_c35070, without toxin (residues 1-1875) and full length plus antitoxin WP_144084685). The DNA sequence encoding for the respective protein was cloned from genomic DNA into vector pTSL (a homemade modified pET-22a vector as described in Taylor et al. (2016)), encoding an N-terminal StrepII-SlyD-TEV tag (resulting in the additional N-terminal residues GSGS after TEV cleavage). The protein was overexpressed in *E. coli* B834 (DE3) (a methionine auxotroph). The cultures were grown to an OD of 0.6 in the presence of 100 μg/ml ampicillin, cooled down to 18°C and induced with 1 mM IPTG. At this point, another 100 μg/ml of ampicillin was added. The cultures were grown overnight at 18°C. The following day, the cells were harvested by centrifugation 5180× g for 20 min at 4°C and flash frozen. Frozen cells (∼30 g) were resuspended in 100 ml of lysis/wash buffer (100 mM TRIS-HCl, 150 mM NaCl, pH 8.0) supplemented with one SIGMAFAST protease inhibitor tablet. Cells were lysed by sonication (15 min, 8 sec on, 12 sec off, T<= 12°C, 60 % amplitude) on ice.

The sample was then purified using StrepTactin Macroprep resin (5 ml) by gravity flow. After loading of the lysate, the resin was washed with 5 column volumes of lysis/wash buffer. The protein was eluted with 0.8, 0.7, 0.7 and 0.8 column volumes of elution buffer (lysis/wash buffer supplemented with 2.5 mM desthiobiotin). The first two fractions were pooled, and digested overnight with 1:1,000 mass:mass of TEV at 4°C while dialysing in TEV buffer (20 mM TRIS, 100 mM NaCl, 2mM β- mercaptoethanol, pH 8.0). The next day, the protein solution (∼20 ml) was diluted two-fold with MonoQ buffer A (20 mM TRIS pH 8.0). The protein was centrifuged for 20 min at 34,957×g and the supernatant was loaded on a GE MonoQ 10/100 GL column. After elution with a 0-500 mM NaCl gradient, fractions containing protein were pooled and concentrated on Vivaspin concentrator with a 50,000 MW cut-off PES membrane, followed by injection on a GE Superdex HiLoad 16/60 column. Peak fractions were concentrated as described above, and the sample was used for crystallisation trials.

### Cloning, expression and purification of S. bongori Rhs+toxin

A. *S. bongori* N268-08 protein (uniprot S5MXP0, ORF A464_2174) (residues 360-1530) was cloned into a modified pBAD33 vector that resembled the pTSL vector described above as previously described (Taylor et al. 2016). The corresponding antitoxin (ORF A464_2173) was cloned into the vector pBAV1K-T5-gfp. The vector pBAV1K-T5-gfp was a gift from Ichiro Matsumura (Addgene plasmid # 26702 ; http://n2t.net/addgene:26702 ; RRID:Addgene_26702) (Bryksin and Matsumura 2010). The culture was grown overnight, and overexpression induced with 2 ml/l of 20% arabinose. The cells were harvested by centrifugation 5180× g for 20 min at 4°C and flash frozen. Frozen cells (∼30 g) were resuspended in 100 ml of lysis/wash buffer (100 mM TRIS-HCl, 150 mM NaCl, pH 8.0) supplemented with one SIGMAFAST protease inhibitor tablet. Cells were lysed by sonication (15 min, 8 sec on, 12 sec off, T<= 12°C, 60 % amplitude) on ice. The purification protocol was the same as for the other proteins.

### Crystallisation

#### A. S. bongori Rhs core domain crystallisation

Protein was diluted to 3 mg/ml using MonoQ buffer A and 1 μl was mixed with 2 μl of reservoir solution (0.1 M MES pH 7, 20% PEE 270, 3.67% acetone) in a hanging drop vapor diffusion experiment at 19°C to obtain the crystals for the native dataset used for SIRAS phasing and for refinement. Before the diffraction experiment, crystals were cryoprotected by soaking in a solution consisting of 0.1M MES pH7 and 35% PEE270.

For the Hg(II) derivative dataset used in SIRAS phasing, the crystal was grown in a similar fashion but using a reservoir solution comprising 0.1 M HEPES pH 7, 20% PEE 270. Derivatisation was performed by soaking the crystal for a couple of minutes in reservoir solution supplemented with 12.5 mM thimerosal. Crystals were back soaked and cryoprotected in mother liquor with an increased PEE 270 concentration (35%).

#### A. S. bongori Rhs core domain (extended N-terminus) crystallisation

Protein solution (1 μl, 4 mg/ml) was mixed with 1 μl reservoir solution (0.1 M MES pH 6.5, 30% ethylene glycol, 7% PEG 8,000) and 0.2 μl of JBScreen Plus compound B9 (2M sodium fluoride) in a hanging drop vapor diffusion experiment at 19°C. A crystal was recovered and flash frozen without the addition of additional cryoprotectant.

#### A. S. bongori *Rhs + toxin*

A hanging drop vapor diffusion experiment was set up at 19°C in which 1 μl of protein solution (2.29 g/l) was mixed with 1 μl reservoir solution (0.1 M MES pH 7, 2% ethylene glycol, 7% PEG 8,000). A crystal was flash frozen without additional cryoprotectant.

#### A. mimigardefordensis DPN7T Rhs crystallisation

Protein solution (1 μl, 13.8 mg/ml) was mixed with 1 μl of mother liquor (0.1 M TRIS pH 8.5, 45% PEP426, 400 mM KCl) for a hanging drop vapor diffusion experiment at 19°C. The crystal was directly flash frozen in liquid nitrogen.

### Data collection

X-ray diffraction data from the four crystals was collected at a wavelength of 1.0000 Å at the PXI beamline (PXII beamline for N-terminal extension of *S. bongori* Rhs core at a wavelength of 0.9788 Å) of the Swiss Light Source of the Paul Scherrer Institute (Switzerland).

### Structure solution and refinement

A. S. bongori *Rhs core domain*

The structure was solved with SIRAS using autoSHARP. The phases were further improved with SHARP, before automatic model building with ARP/wARP. The phases from ARP/wARP were used as input for buccaneer. This model was further rebuilt with Coot, before refinement against the native data from the SIRAS experiment with PHENIX. Subsequently, we used a dataset diffracting to high resolution (with R-free flags transferred with CCP4 uniquify) for refinement with PHENIX after rebuilding, followed by refinement with autoBUSTER and PHENIX.

#### A. S. bongori Rhs core domain (extended N-terminal region)

The structure was solved by molecular replacement with PHASER, using an ensemble of the two molecules in the asymmetric of the Rhs core domain structure without N-terminal extension. R-free flags were transferred from the *S. bongori* Rhs core domain native dataset because these crystals are isomorphous. The structure was rebuilt with Buccaneer and Coot, and refined with autoBUSTER and PHENIX.

#### A. S. bongori *Rhs + toxin*

This structure was solved by molecular replacement with Rhs without N-terminal extension as a search model. The model was rebuilt in Coot and refined using autoBUSTER and PHENIX.

#### A. mimigardefordensis *DPN7T Rhs*

The structure was solved using molecular replacement with an ensemble of the two molecules in the asymmetric of the Rhs core domain structure without N-terminal extension. This was followed by model rebuilding in Coot, and refinement with autoBUSTER and PHENIX. The comparison structure of VgrG-linked Rhs from *A. mimigardefordensis* was predicted using AlphaFold v 2.1.1 (Jumper et al. 2021).

### Bioinformatics: T6SS effector analysis

T6SS effector sequences were compiled from the SecReT6 v3 T6SS database (76,048 effector sequences) (Li et al. 2015), from a recent publication (Table S2 from Liu et al. (2020), mapping identifiers to 531 sequences in Uniprot and 827 sequences in Uniparc, 99 unidentified identifiers could not be mapped), as well as from InterPro (108 VgrG-Rhs, 2,603 PAAR-Rhs), as displayed in Figure 4 A.

Duplicates by sequence were removed using Seqkit (Shen et al. 2016) (18,330 sequences left) and domains annotated using the Conserved Domains and Protein Classification (Marchler-Bauer et al. 2011, Lu et al. 2020). Results were obtained for 17,552 sequences, the expect value threshold was 0.01 and the output mode was concise. Prefix categories (no prefix, PAAR, VgrG, Hcp, VipA, Rhs, PAAR-Rhs and VgrG-Rhs) and effector categories were mapped to CDD hits according to Table S4 and S5, respectively, using Python pandas v2.0.1 (McKinney and Others 2010). All CDD results with annotated domains can be found in Table S3.

Effectors for which an annotation was found using CDD were then plotted for prefix and effector categories in bar charts. These effectors were analysed for their molecular weight and pI using Biopython and SeqIO (Cock et al. 2009). In silico cleavage of Rhs effectors was performed at the canonical PxxxxDPxG[LWF], or, if this motif was not present, at the shorter DPxG[LWF] motif. If neither motif was present, the protein was not analysed. If two cleavage sites were present, the shorter fragment was kept, in accordance with the CDD annotation of toxin domains. Seqkit (Shen et al. 2016) was used for the in silico cleavage of Rhs effectors and the sequence handling. The sequence of the C-terminal cleavage product was extracted and analysed. As comparison, the annotated effector domains for PAAR- and VgrG-linked effectors were extracted, extended to the C-terminus and analysed. Rhs effectors were also analysed for the presence of TMDs using TMHMM (Krogh et al. 2001) and DeepTMHMM (Hallgren et al. 2022). Rhs effectors were in silico cleaved as described above and the sequence of the N-terminal cleavage product was extracted and analysed for TMDs, since the algorithms otherwise detect the C-terminal cleavage site as TMD and miss a potential N-terminal TMD in many cases.

To search for sequences similar to *Vibrio parahaemolyticus* VIR, the VIR sequence was used in jackhmmer to find more sequences, accepting the results after two iterations, since in later iterations, all new hits were non-Rhs proteins. After creating a hidden Markov model from the jackhmmer hits, hmmsearch was used to search all Rhs sequences for which we could identify a toxin domain and all T6SS effector sequences in our list as comparison.

### Structure prediction of Salmonella bongori PAAR-Rhs toxin domain and incomplete Rhs proteins

The toxin domain of *Salmonella bongori* PAAR-Rhs (residues 1310 - 1420) was used for structure prediction with AlphaFold2 v 2.1.1. Structure predictions were manually inspected and structural homologies were searched using DALI PDB search (Holm 2022). Structure prediction for incomplete Rhs cores was performed using AlphaFold2 v 2.1.1.

### Data visualisation and figures

UCSF Chimera and coot were used to visualise molecular models and maps. GraphPad Prism 9 was used for plotting and statistical analysis. Maps and models were superimposed for comparison using Coot’s map transformation by LSQ model fit feature. All electrostatic potentials were calculated using the APBS web service (Jurrus et al. 2018). Figures were created using Adobe Illustrator.

### Data availability

Molecular models for the *S. bongori* Rhs core, the *S. bongori* Rhs core with extended N-terminus and *S. bongori* Rhs core + toxin have been submitted to the PDB under the accession codes 8QJD, 8QJB and 8QJC, respectively. The *A. mimigardefordensis* VgrG-Rhs model has been submitted to the PDB under the accession code 8QJA.

Supplementary tables and files:

ST1: Refinement statistics

ST2: Significant DALI hits against PDB100 for AlphaFold prediction of *S. bongori* PAAR-linked Rhs toxin domain.

ST3: 34,151 CDD hits for 18,330 unique T6SS effector sequences.

ST4: Annotations for prefix domains and Rhs (Hcp, PAAR, Rhs, VgrG, VipA) for CDD results.

ST5: Annotations for effector categories for CDD results. ST6: jackhmmer results for *Vibrio parahaemolyticus* VIR ST7: VIR Hidden Markov model

ST8: hmmsearch hits for VIR hidden Markov model

### Other supplementary data

Fasta-file with all T6SS effectors, duplicates removed

## Supporting information

Refinement statistics

Significant DALI hits against PDB100 for AlphaFold prediction of S. bongori PAAR-linked Rhs toxin domain

34,151 CDD hits for 18,330 unique T6SS effector sequences

Annotations for prefix domains and Rhs (Hcp, PAAR, Rhs, VgrG, VipA) for CDD results

Annotations for effector categories for CDD results

jackhmmer results for Vibrio parahaemolyticus VIR

VIR Hidden Markov model

hmmsearch hits for VIR hidden Markov model

## Acknowledgements

We acknowledge the Paul Scherrer Institut, Villigen, Switzerland for provision of synchrotron radiation beamtime at beamlines PXI and PXII of the SLS. We thank Yumeng Yan and Victor Klein De Sousa for help with TMD analyses and bioinformatics. The project was partly funded by the Swiss National Science Foundation grant 310030_144243 to P.G.L. The Novo Nordisk Foundation Center for Protein Research is supported financially by the Novo Nordisk Foundation (NNF14CC0001). N.M.I.T. acknowledges support from a DFF grant (8123-00002B), NNF Hallas-Møller Emerging Investigator grant (NNF17OC0031006), and the EMBO Young Investigator programme.

## Author contributions

C.S.K performed data analysis, designed and performed the bioinformatics analyses, and wrote the paper. M.M.S. designed and performed experiments. P.G.L designed experiments and wrote the paper. N.M.I.T designed experiments, performed experiments and data analysis, and wrote the paper.

## Primers

*Salmonella bongori* A464_2174:

360-1420:

FW: 5’ CCAGGGCAGCGGATCCGAGCCGGTGGACATCGGAACC 3’

RV: 5’ GGTGGTGGTGCTCGAGTTATAGCCCAAGCGGATCGATC 3’

348-1420:

FW: 5’ AATAGGATCCGGGGATGGTGTAAACAGTATG 3’

RV: 5’ GGTGGTGGTGCTCGAGTTATAGCCCAAGCGGATCGATC 3’

360-1530:

FW: 5’ CCAAGCTTAAGCTTGTCGACTCATTGTGTTGTTAACCTTCGG 3’

RV: 5’ GGTGGTGGTGCTCGAGTTATAGCCCAAGCGGATCGATC 3’

*Salmonella bongori* A464_2173 (antitoxin):

FW: 5’ AGAGGAGAAATACTGATGAACATTGATTTCTCAGTAAAA 3’

RV: 5’ GCCGCTACTAGTATATTAGAGCTTATTTATAGCCT 3’

A. mimigardefordensis AHG65567: 1-1875:

FW: 5’ CCAGGGCAGGATCCAACCGTACCGTAAATGCCAC 3’

RV: 5’ GTGCGGCCGCAAGCTTCAATCCGAAAGGATCAATCCA 3’

---

**Figure S1:**
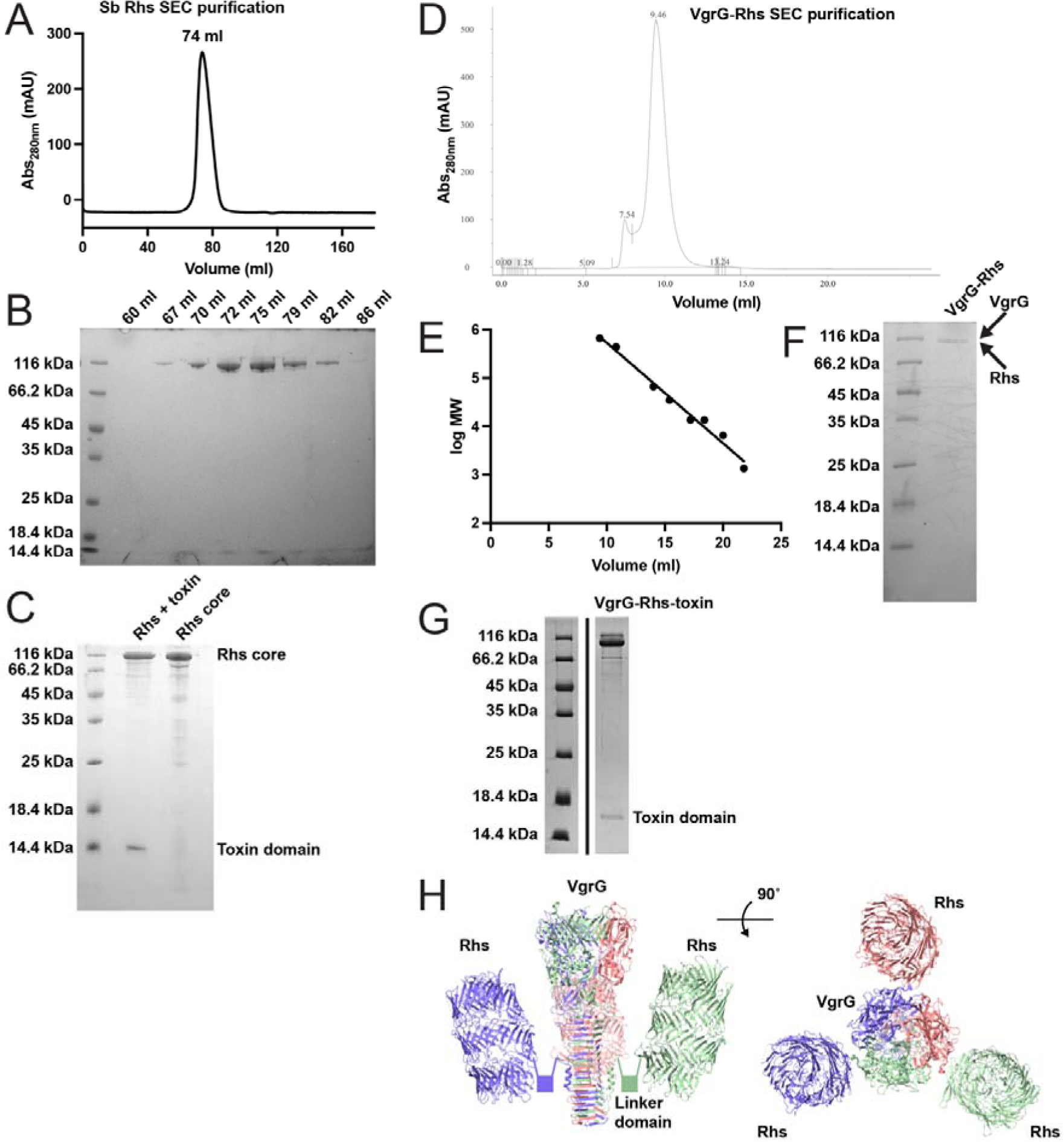
Purification of Rhs proteins. A) SEC trace of *S. bongori* PAAR-linked Rhs core domain. The Rhs core elutes at 74 ml on a Superdex 200 HiLoad 16/600 column. B) Coomassie SDS-PAGE of fractions of *S. bongori* PAAR-linked Rhs core SEC run in A). C) Coomassie SDS- PAGE of *S. bongori* PAAR-linked Rhs + toxin in comparison with the Rhs core domain only. A clear band is visible for the autocleaved toxin domain in the Rhs + toxin sample. D) SEC trace of *A. mimigardefordensis* VgrG-linked Rhs. VgrG-Rhs elutes at 9.46 ml on a Superdex 200 10/300 column. E) Calibration curve of the Superdex 200 10/300 column. F) Coomassie SDS- PAGE of purified *A. mimigardefordensis* VgrG-Rhs, showing the protein is cleaved into the VgrG and Rhs domains. G) SDS-PAGE of purified full-length AHG65567. A band runs at ∼16 kDa, corresponding with the cleaved C-terminus (residues 1876-2012, 15.9 kDa theoretical molecular weight). H) Schematic of Rhs containers that are genetically linked to VgrG, using the *A. mimigardefordensis* Rhs container structure determined here and a VgrG structure from *Pseudomonas aeruginosa* (PDB ID 4mtk). For a spike of trimeric VgrG, three Rhs containers are attached.

**Figure S2:**
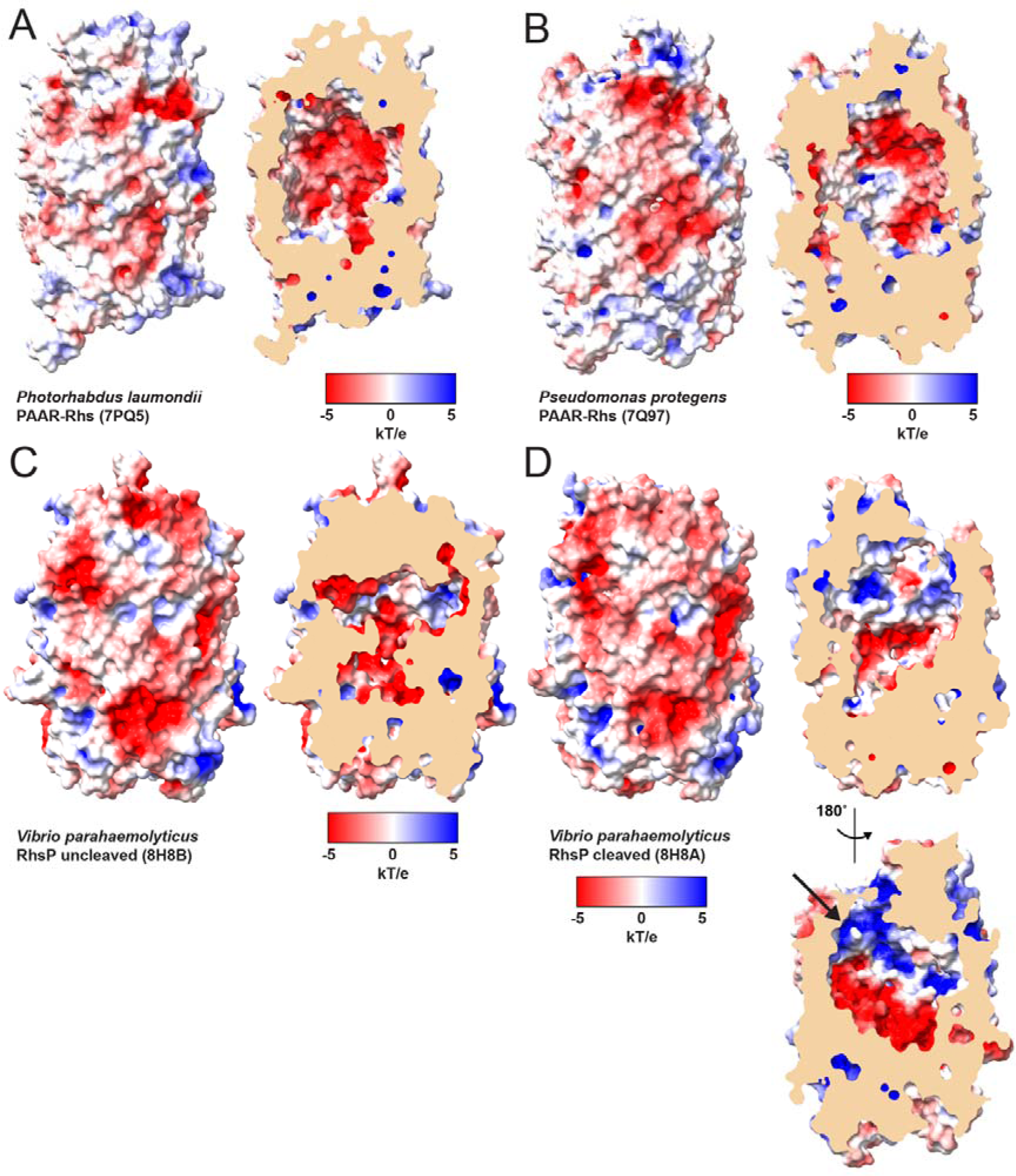
Electrostatic surface of recently published Rhs structures. A – D) Surface representation of the recently published Rhs structures PAAR-Rhs from *Photorhabdus laumondii* (7PQ5), PAAR-Rhs from *Pseudomonas protegens* (7Q97) and uncleaved and cleaved RhsP from *Vibrio parahaemolytics* (8H8B and 8H8A, respectively). Surfaces are coloured according to electrostatic potential, as calculated using APBS, in a side view and clipped to expose the negative container inside. The positively charged patch close to the hypothesised exit crevice in cleaved RhsP is indicated with an arrow in D).

**Figure S3:**
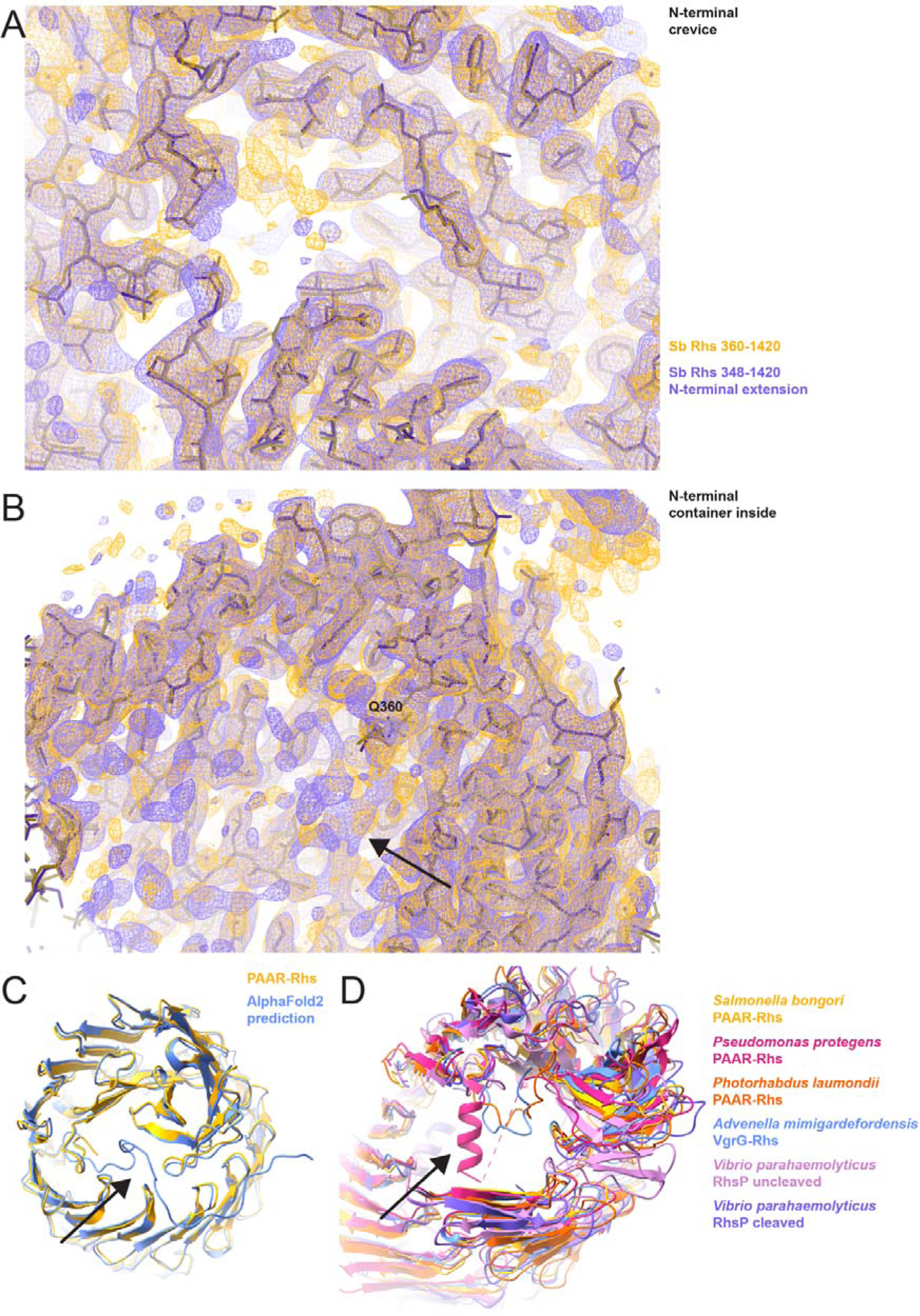
Rhs plug domain. A) Overlay of *S. bongori* Rhs core (yellow) and N-terminally extended Rhs map (purple) and molecular model, showing the N-terminal crevice. No additional density plugging the crevice can be detected in the extended Rhs. B) Overlay of *S*. *bongori* Rhs core (yellow) and N-terminally extended Rhs map (purple) and molecular model, showing the N-terminus and the inside of the container. Although no additional density preceding from Glu-360 can be traced in the extended Rhs, some weak density is present inside the container (indicated with arrow). C) Overlay of *S. bongori* Rhs core (yellow) with an AlphaFold2 prediction (blue), showing that in the prediction, the N-terminus of the *S. bongori* Rhs core closes off the container (arrow). D) Comparison of Rhs container structures from *S. bongori* (this publication, PAAR-Rhs, yellow), *Pseudomonas protegens* (pdb 7Q97, PAAR-Rhs, magenta), *Photorhabdus laumondii* (pdb 7PQ5, PAAR-Rhs, orange) and *A. mimigardefordensis* (this publication, VgrG-Rhs, blue). PAAR-Rhs from *Pseudomonas protegens* is the only Rhs core that carries a so-called anchor helix (arrow). Obstructing residues were hidden from view to display the anchor helix region. The contour level in A and B is 0.7 r.m.s.d.

**Figure S4:**
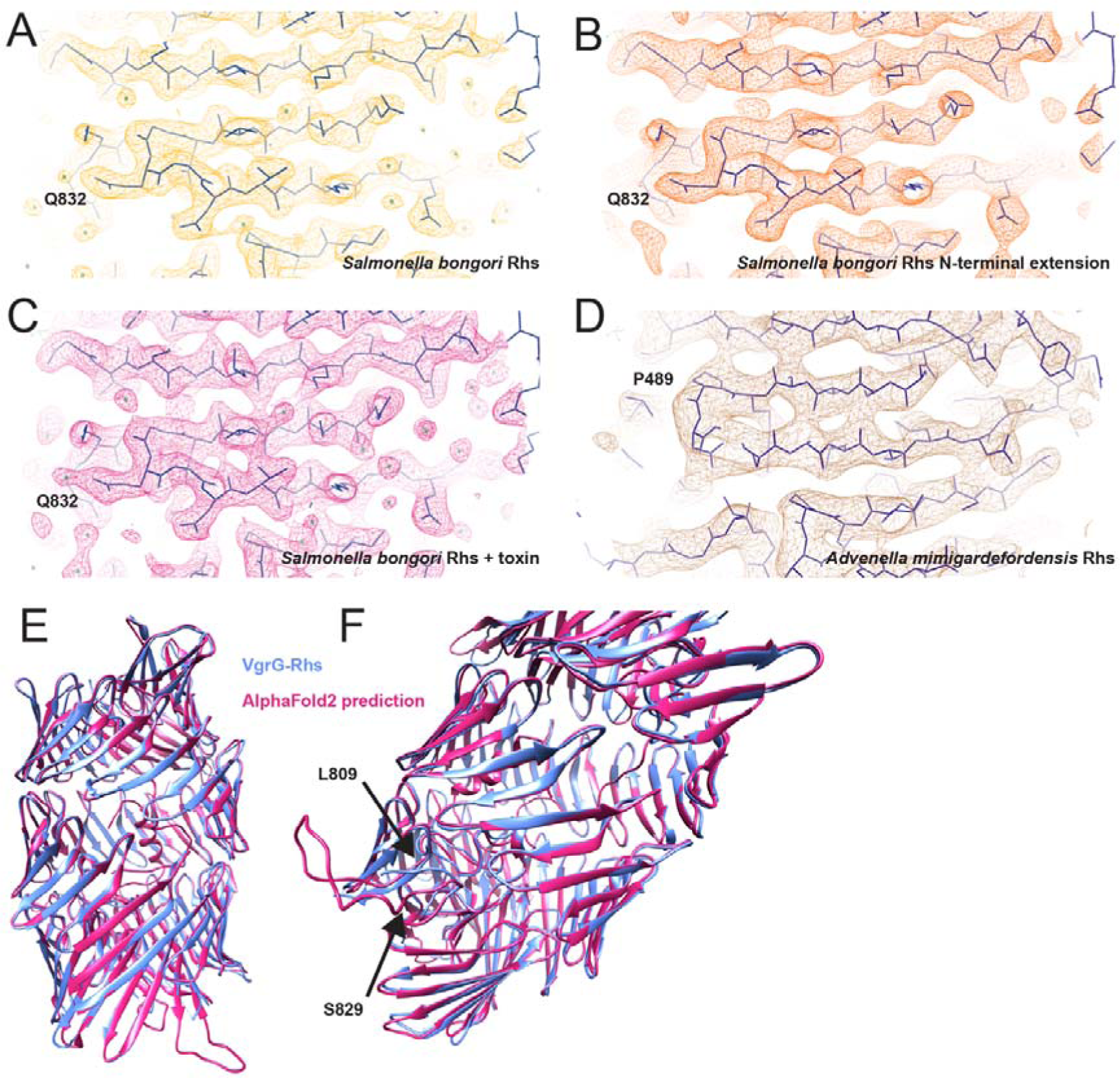
A) − D) Overview over map resolution for A) *S. bongori* PAAR-linked Rhs, B) Rhs N- terminal extension, C) Rhs + toxin and D) *A. mimigardefordensis* VgrG-linked Rhs. E) An AlphaFold2 prediction of VgrG-linked Rhs core from *A. mimigardefordensis* (magenta) agrees very well with the molecular model built into the experimental structure (blue). F) In the area of residues 809-829, however, AlphaFold2 predicts one big loop instead of a β-strand.

**Figure S5:**
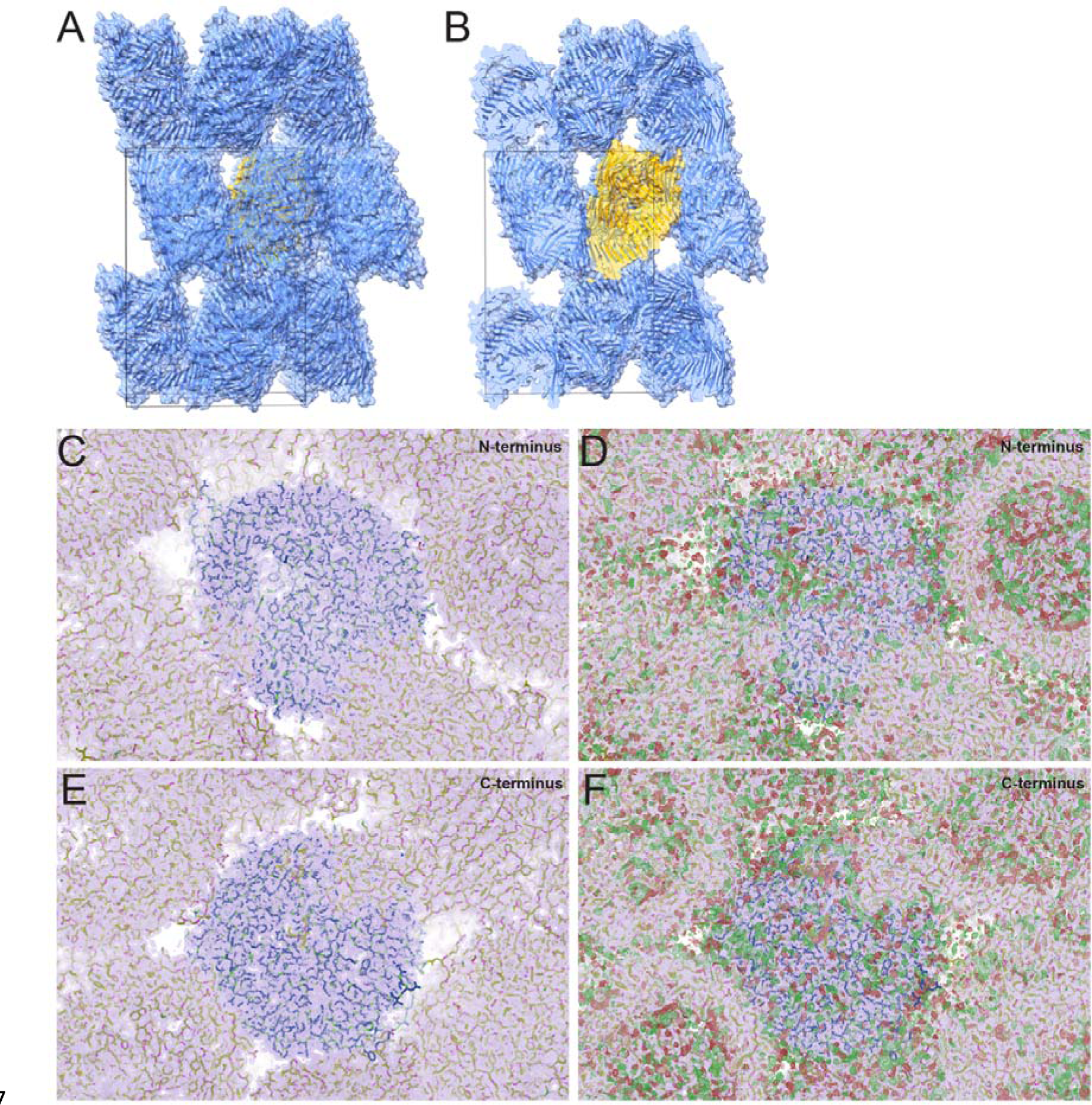
Crystal packing of *S. bongori* PAAR-Rhs core + toxin shows no large unassigned density outside of the Rhs core. A) Model of the Rhs core + toxin in the asymmetric unit (yellow), surrounded by its crystallographic symmetry equivalents (blue), in cartoon and surface representation. In B), the models are clipped to reveal some empty space between the symmetry copies. C) 2Fo-Fc and D) 2Fo-Fc and the difference map with built models of the Rhs core + toxin, shown looking onto the N-terminus of the model in the asymmetric unit (centre). There is no large unassigned density present outside of the Rhs core that could account for the toxin domain. E) 2Fo-Fc and F) 2Fo-Fc and the difference map with built models of the Rhs core + toxin, shown looking onto the C- terminus of the model in the asymmetric unit (centre). There is no large unassigned density present outside of the Rhs core that could account for the toxin domain. The contour levels in C − F are 1 r.m.s.d and 2.2 r.m.s.d for the 2Fo-Fc and the difference map, respectively.

**Figure S6:**
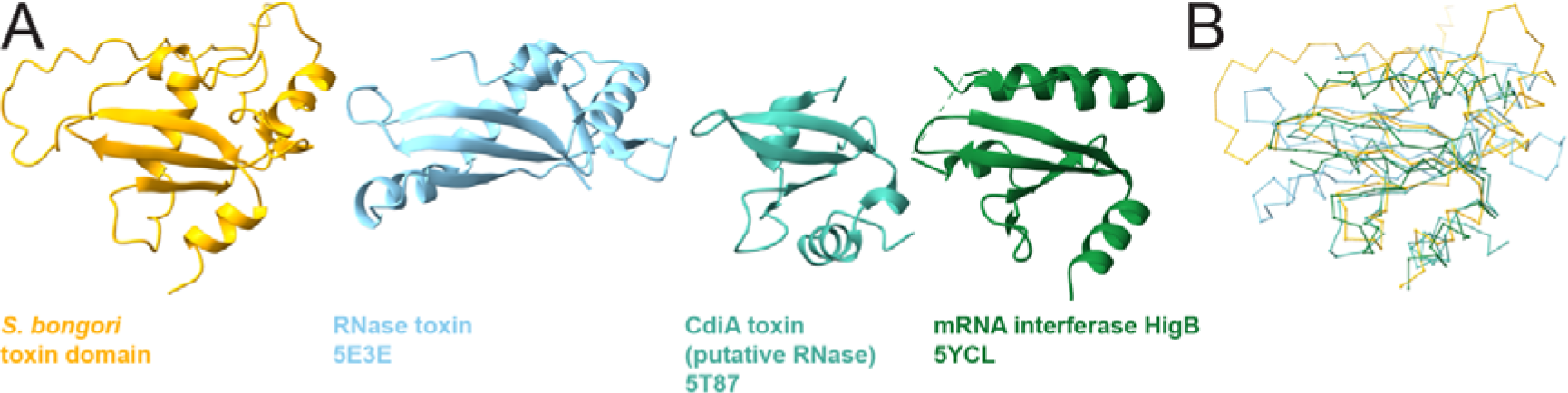
The *S. bongori* Rhs toxin domain has an RNase fold. A) Structural comparison of the *S. bongori* Rhs toxin domain with the three top matches in the DALI search against the PDB. B) Overlay of C-alpha chains of the *S. bongori* Rhs toxin domain with the three top matches in the DALI search against the PDB.

**Figure S7:**
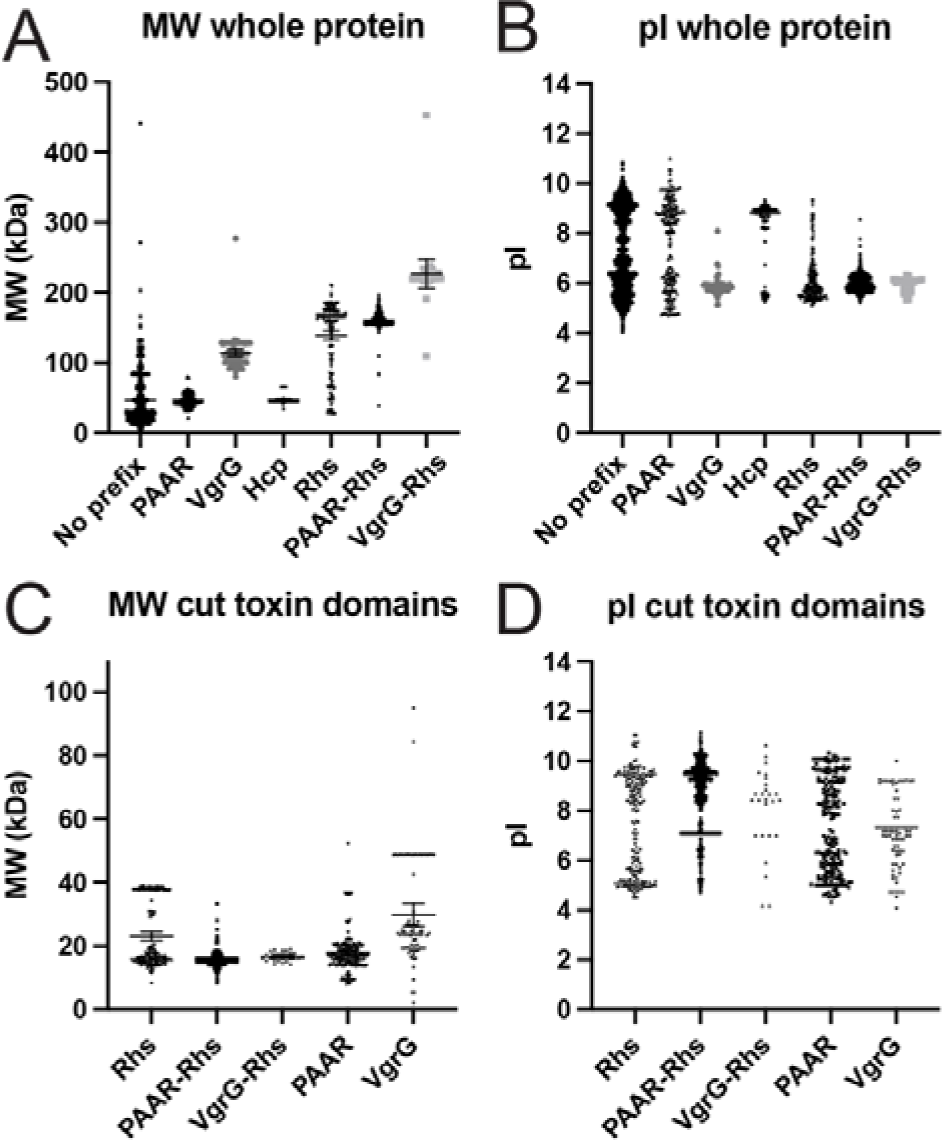
Molecular weight and isoelectric point distributions of Rhs and non-Rhs effectors. A) Molecular weights for whole protein effectors with annotated effector domain. B) Isoelectric points for whole protein effectors with annotated effector domain. C) Molecular weights for in silico cleaved Rhs effectors and C-terminal effector domains for PAAR- and VgrG-linked effectors. D) Isoelectric points for in silico cleaved Rhs effectors and C-terminal effector domains for PAAR- and VgrG-linked effectors. For molecular weights, the mean and the 95% confidence interval are shown.

**Figure S8:**
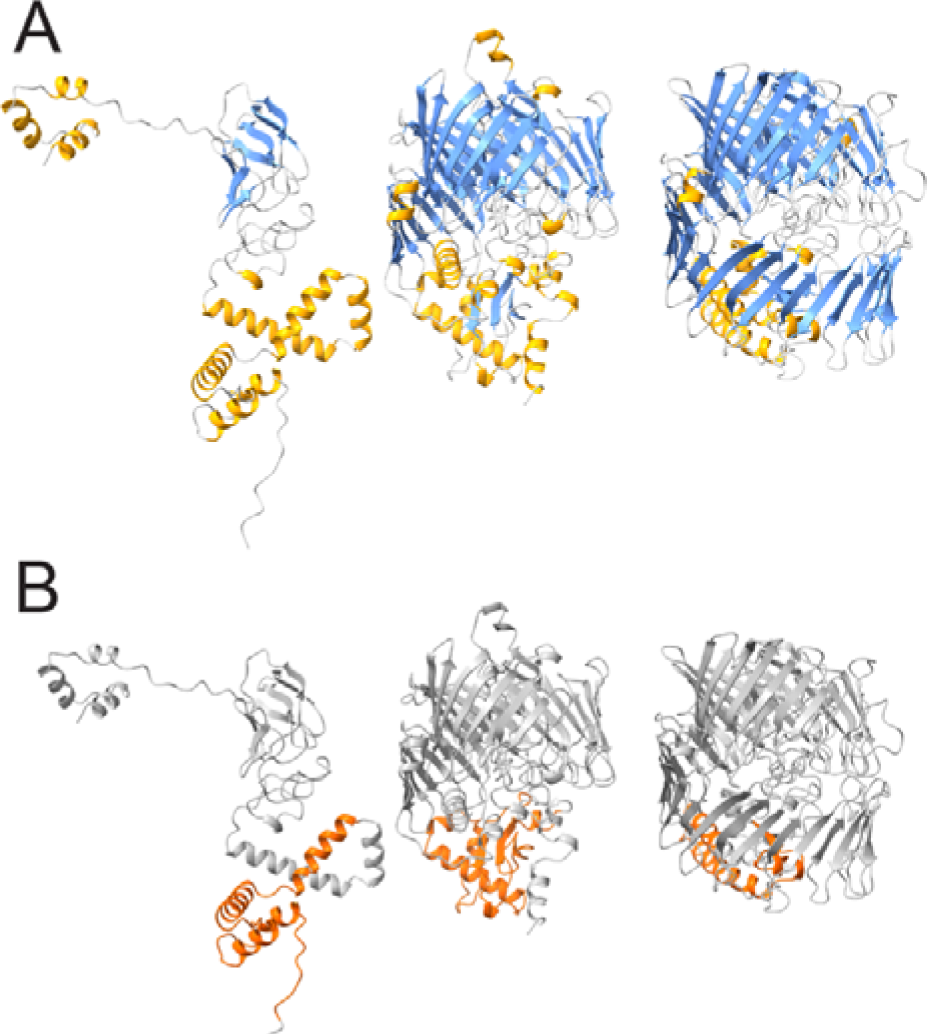
Three examples of AlphaFold2 structure predictions of small Rhs proteins in our collection, showing that they constitute an incomplete Rhs container, A) coloured according to secondary structure and B) the toxin domain coloured in orange.

## References

Alcoforado Diniz, J., and S. J. Coulthurst. 2015. “Intraspecies Competition in Serratia marcescens Is Mediated by Type VI-Secreted Rhs Effectors and a Conserved Effector-Associated Accessory Protein.” J Bacteriol 197 (14):2350–60. doi: 10.1128/JB.00199-15.

Alcoforado Diniz, J., Y. C. Liu, and S. J. Coulthurst. 2015. “Molecular weaponry: diverse effectors delivered by the Type VI secretion system.” Cell Microbiol 17 (12):1742–51. doi: 10.1111/cmi.12532.

Basler, M., M. Pilhofer, G. P. Henderson, G. J. Jensen, and J. J. Mekalanos. 2012. “Type VI secretion requires a dynamic contractile phage tail-like structure.” Nature 483 (7388):182–6. doi: 10.1038/nature10846.

Batot, G., K. Michalska, G. Ekberg, E. M. Irimpan, G. Joachimiak, R. Jedrzejczak, G. Babnigg, C. S. Hayes, A. Joachimiak, and C. W. Goulding. 2017. “The CDI toxin of Yersinia kristensenii is a novel bacterial member of the RNase A superfamily.” Nucleic Acids Res 45 (9):5013–5025. doi: 10.1093/nar/gkx230.

Bonemann, G., A. Pietrosiuk, A. Diemand, H. Zentgraf, and A. Mogk. 2009. “Remodelling of VipA/VipB tubules by ClpV-mediated threading is crucial for type VI protein secretion.” EMBO J 28 (4):315–25. doi: 10.1038/emboj.2008.269.

Bryksin, A. V., and I. Matsumura. 2010. “Rational design of a plasmid origin that replicates efficiently in both gram-positive and gram-negative bacteria.” PLoS One 5 (10):e13244. doi: 10.1371/journal.pone.0013244.

Burkinshaw, B. J., X. Liang, M. Wong, A. N. H. Le, L. Lam, and T. G. Dong. 2018. “A type VI secretion system effector delivery mechanism dependent on PAAR and a chaperone-co-chaperone complex.” Nat Microbiol 3 (5):632–640. doi: 10.1038/s41564-018-0144-4.

Busby, J. N., S. Panjikar, M. J. Landsberg, M. R. Hurst, and J. S. Lott. 2013. “The BC component of ABC toxins is an RHS-repeat-containing protein encapsulation device.” Nature 501 (7468):547–50. doi: 10.1038/nature12465.

Carobbi, A., S. Di Nepi, C. M. Fridman, Y. Dar, R. Ben-Yaakov, I. Barash, D. Salomon, and G. Sessa. 2022. “An antibacterial T6SS in Pantoea agglomerans pv. betae delivers a lysozyme-like effector to antagonize competitors.” Environ Microbiol 24 (10):4787–4802. doi: 10.1111/1462-2920.16100.

Chang, Y. W., L. A. Rettberg, D. R. Ortega, and G. J. Jensen. 2017. “In vivo structures of an intact type VI secretion system revealed by electron cryotomography.” EMBO Rep 18 (7):1090–1099. doi: 10.15252/embr.201744072.

Cock, P. J., T. Antao, J. T. Chang, B. A. Chapman, C. J. Cox, A. Dalke, I. Friedberg, T. Hamelryck, F. Kauff, B. Wilczynski, and M. J. de Hoon. 2009. “Biopython: freely available Python tools for computational molecular biology and bioinformatics.” Bioinformatics 25 (11):1422–3. doi: 10.1093/bioinformatics/btp163.

Coulthurst, S. 2019. “The Type VI secretion system: a versatile bacterial weapon.” Microbiology (Reading) 165 (5):503–515. doi: 10.1099/mic.0.000789.

De Maayer, P., S. N. Venter, T. Kamber, B. Duffy, T. A. Coutinho, and T. H. Smits. 2011. “Comparative genomics of the Type VI secretion systems of Pantoea and Erwinia species reveals the presence of putative effector islands that may be translocated by the VgrG and Hcp proteins.” BMC Genomics 12:576. doi: 10.1186/1471-2164-12-576.

Donato, S. L., C. M. Beck, F. Garza-Sanchez, S. J. Jensen, Z. C. Ruhe, D. A. Cunningham, I. Singleton, D. A. Low, and C. S. Hayes. 2020. “The beta- encapsulation cage of rearrangement hotspot (Rhs) effectors is required for type VI secretion.” Proc Natl Acad Sci U S A. doi: 10.1073/pnas.1919350117.

Durand, E., C. Cambillau, E. Cascales, and L. Journet. 2014. “VgrG, Tae, Tle, and beyond: the versatile arsenal of Type VI secretion effectors.” Trends Microbiol 22 (9):498–507. doi: 10.1016/j.tim.2014.06.004.

Gatsogiannis, C., F. Merino, D. Roderer, D. Balchin, E. Schubert, A. Kuhlee, M. Hayer-Hartl, and S. Raunser. 2018. “Tc toxin activation requires unfolding and refolding of a beta-propeller.” Nature 563 (7730):209–213. doi: 10.1038/s41586-018-0556-6.

Green, E. R., and J. Mecsas. 2016. “Bacterial Secretion Systems: An Overview.” Microbiol Spectr 4 (1). doi: 10.1128/microbiolspec.VMBF-0012-2015.

Gunther, P., D. Quentin, S. Ahmad, K. Sachar, C. Gatsogiannis, J. C. Whitney, and S. Raunser. 2022. “Structure of a bacterial Rhs effector exported by the type VI secretion system.” PLoS Pathog 18 (1):e1010182. doi: 10.1371/journal.ppat.1010182.

Hachani, A., T. E. Wood, and A. Filloux. 2016. “Type VI secretion and anti-host effectors.” Curr Opin Microbiol 29:81–93. doi: 10.1016/j.mib.2015.11.006.

Hallgren, Jeppe, Konstantinos D. Tsirigos, Mads Damgaard Pedersen, José Juan Almagro Armenteros, Paolo Marcatili, Henrik Nielsen, Anders Krogh, and Ole Winther. 2022. “DeepTMHMM predicts alpha and beta transmembrane proteins using deep neural networks.” *bioRxiv*:2022.04.08.487609. doi: 10.1101/2022.04.08.487609.

Holm, L. 2022. “Dali server: structural unification of protein families.” Nucleic Acids Res. doi: 10.1093/nar/gkac387.

Hood, R. D., P. Singh, F. Hsu, T. Guvener, M. A. Carl, R. R. Trinidad, J. M. Silverman, B. B. Ohlson, K. G. Hicks, R. L. Plemel, M. Li, S. Schwarz, W. Y. Wang, A. J. Merz, D. R. Goodlett, and J. D. Mougous. 2010. “A type VI secretion system of Pseudomonas aeruginosa targets a toxin to bacteria.” Cell Host Microbe 7 (1):25–37. doi: 10.1016/j.chom.2009.12.007.

Jackson, A. P., G. H. Thomas, J. Parkhill, and N. R. Thomson. 2009. “Evolutionary diversification of an ancient gene family (rhs) through C-terminal displacement.” BMC Genomics 10:584. doi: 10.1186/1471-2164-10-584.

Jumper, J., R. Evans, A. Pritzel, T. Green, M. Figurnov, O. Ronneberger, K. Tunyasuvunakool, R. Bates, A. Zidek, A. Potapenko, A. Bridgland, C. Meyer, S. A. A. Kohl, A. J. Ballard, A. Cowie, B. Romera-Paredes, S. Nikolov, R. Jain, J. Adler, T. Back, S. Petersen, D. Reiman, E. Clancy, M. Zielinski, M. Steinegger, M. Pacholska, T. Berghammer, S. Bodenstein, D. Silver, O. Vinyals, A. W. Senior, K. Kavukcuoglu, P. Kohli, and D. Hassabis. 2021. “Highly accurate protein structure prediction with AlphaFold.” Nature 596 (7873):583–589. doi: 10.1038/s41586-021-03819-2.

Jurenas, D., N. Fraikin, F. Goormaghtigh, and L. Van Melderen. 2022. “Biology and evolution of bacterial toxin-antitoxin systems.” Nat Rev Microbiol. doi: 10.1038/s41579-021-00661-1.

Jurenas, D., L. T. Rosa, M. Rey, J. Chamot-Rooke, R. Fronzes, and E. Cascales. 2021. “Mounting, structure and autocleavage of a type VI secretion- associated Rhs polymorphic toxin.” Nat Commun 12 (1):6998. doi: 10.1038/s41467-021-27388-0.

Jurrus, E., D. Engel, K. Star, K. Monson, J. Brandi, L. E. Felberg, D. H. Brookes, L. Wilson, J. Chen, K. Liles, M. Chun, P. Li, D. W. Gohara, T. Dolinsky, R. Konecny, D. R. Koes, J. E. Nielsen, T. Head-Gordon, W. Geng, R. Krasny, G. W. Wei, M. J. Holst, J. A. McCammon, and N. A. Baker. 2018. “Improvements to the APBS biomolecular solvation software suite.” Protein Sci 27 (1):112–128. doi: 10.1002/pro.3280.

Kiraga, J., P. Mackiewicz, D. Mackiewicz, M. Kowalczuk, P. Biecek, N. Polak, K. Smolarczyk, M. R. Dudek, and S. Cebrat. 2007. “The relationships between the isoelectric point and: length of proteins, taxonomy and ecology of organisms.” BMC Genomics 8:163. doi: 10.1186/1471-2164-8-163.

Koskiniemi, S., J. G. Lamoureux, K. C. Nikolakakis, C. T’Kint de Roodenbeke, M. D. Kaplan, D. A. Low, and C. S. Hayes. 2013. “Rhs proteins from diverse bacteria mediate intercellular competition.” Proc Natl Acad Sci U S A 110 (17):7032–7. doi: 10.1073/pnas.1300627110.

Krogh, A., B. Larsson, G. von Heijne, and E. L. Sonnhammer. 2001. “Predicting transmembrane protein topology with a hidden Markov model: application to complete genomes.” J Mol Biol 305 (3):567–80. doi: 10.1006/jmbi.2000.4315.

Leiman, P. G., M. Basler, U. A. Ramagopal, J. B. Bonanno, J. M. Sauder, S. Pukatzki, S. K. Burley, S. C. Almo, and J. J. Mekalanos. 2009. “Type VI secretion apparatus and phage tail-associated protein complexes share a common evolutionary origin.” Proc Natl Acad Sci U S A 106 (11):4154–9. doi: 10.1073/pnas.0813360106.

Li, J., Y. Yao, H. H. Xu, L. Hao, Z. Deng, K. Rajakumar, and H. Y. Ou. 2015. “SecReT6: a web-based resource for type VI secretion systems found in bacteria.” Environ Microbiol 17 (7):2196–202. doi: 10.1111/1462-2920.12794.

Liang, X., R. Moore, M. Wilton, M. J. Wong, L. Lam, and T. G. Dong. 2015. “Identification of divergent type VI secretion effectors using a conserved chaperone domain.” Proc Natl Acad Sci U S A 112 (29):9106–11. doi: 10.1073/pnas.1505317112.

Lien, Y. W., and E. M. Lai. 2017. “Type VI Secretion Effectors: Methodologies and Biology.” Front Cell Infect Microbiol 7:254. doi: 10.3389/fcimb.2017.00254.

Liu, Y., Z. Zhang, F. Wang, D. D. Li, and Y. Z. Li. 2020. “Identification of type VI secretion system toxic effectors using adaptors as markers.” Comput Struct Biotechnol J 18:3723–3733. doi: 10.1016/j.csbj.2020.11.003.

Lu, S., J. Wang, F. Chitsaz, M. K. Derbyshire, R. C. Geer, N. R. Gonzales, M. Gwadz, D. I. Hurwitz, G. H. Marchler, J. S. Song, N. Thanki, R. A. Yamashita, M. Yang, D. Zhang, C. Zheng, C. J. Lanczycki, and A. Marchler-Bauer. 2020. “CDD/SPARCLE: the conserved domain database in 2020.” Nucleic Acids Res 48 (D1):D265–D268. doi: 10.1093/nar/gkz991.

Ma, J., M. Sun, W. Dong, Z. Pan, C. Lu, and H. Yao. 2017. “PAAR-Rhs proteins harbor various C-terminal toxins to diversify the antibacterial pathways of type VI secretion systems.” Environ Microbiol 19 (1):345–360. doi: 10.1111/1462-2920.13621.

Marchler-Bauer, A., S. Lu, J. B. Anderson, F. Chitsaz, M. K. Derbyshire, C. DeWeese-Scott, J. H. Fong, L. Y. Geer, R. C. Geer, N. R. Gonzales, M. Gwadz, D. I. Hurwitz, J. D. Jackson, Z. Ke, C. J. Lanczycki, F. Lu, G. H. Marchler, M. Mullokandov, M. V. Omelchenko, C. L. Robertson, J. S. Song, N. Thanki, R. A. Yamashita, D. Zhang, N. Zhang, C. Zheng, and S. H. Bryant. 2011. “CDD: a Conserved Domain Database for the functional annotation of proteins.” Nucleic Acids Res 39 (Database issue):D225-9. doi: 10.1093/nar/gkq1189.

McKinney, W., and Others. 2010. “Data structures for statistical computing in python.” Proceedings of the 9th Python in Science Conference 445:51--56. doi: 10.25080/Majora-92bf1922-00a.

Meusch, D., C. Gatsogiannis, R. G. Efremov, A. E. Lang, O. Hofnagel, I. R. Vetter, K. Aktories, and S. Raunser. 2014. “Mechanism of Tc toxin action revealed in molecular detail.” Nature 508 (7494):61–5. doi: 10.1038/nature13015.

Pei, T. T., H. Li, X. Liang, Z. H. Wang, G. Liu, L. L. Wu, H. Kim, Z. Xie, M. Yu, S. Lin, P. Xu, and T. G. Dong. 2020. “Intramolecular chaperone-mediated secretion of an Rhs effector toxin by a type VI secretion system.” Nat Commun 11 (1):1865. doi: 10.1038/s41467-020-15774-z.

Pukatzki, S., A. T. Ma, A. T. Revel, D. Sturtevant, and J. J. Mekalanos. 2007. “Type VI secretion system translocates a phage tail spike-like protein into target cells where it cross-links actin.” Proc Natl Acad Sci U S A 104 (39):15508–13. doi: 10.1073/pnas.0706532104.

Pukatzki, S., A. T. Ma, D. Sturtevant, B. Krastins, D. Sarracino, W. C. Nelson, J. F. Heidelberg, and J. J. Mekalanos. 2006. “Identification of a conserved bacterial protein secretion system in Vibrio cholerae using the Dictyostelium host model system.” Proc Natl Acad Sci U S A 103 (5):1528–33. doi: 10.1073/pnas.0510322103.

Quentin, D., S. Ahmad, P. Shanthamoorthy, J. D. Mougous, J. C. Whitney, and S. Raunser. 2018. “Mechanism of loading and translocation of type VI secretion system effector Tse6.” Nat Microbiol 3 (10):1142–1152. doi: 10.1038/s41564-018-0238-z.

Repizo, G. D., M. Espariz, J. L. Seravalle, and S. P. Salcedo. 2019. “Bioinformatic Analysis of the Type VI Secretion System and Its Potential Toxins in the Acinetobacter Genus.” Front Microbiol 10:2519. doi: 10.3389/fmicb.2019.02519.

Ruhe, Z. C., D. A. Low, and C. S. Hayes. 2020. “Polymorphic Toxins and Their Immunity Proteins: Diversity, Evolution, and Mechanisms of Delivery.” Annu Rev Microbiol 74:497–520. doi: 10.1146/annurev-micro-020518-115638.

Shen, W., S. Le, Y. Li, and F. Hu. 2016. “SeqKit: A Cross-Platform and Ultrafast Toolkit for FASTA/Q File Manipulation.” PLoS One 11 (10):e0163962. doi: 10.1371/journal.pone.0163962.

Shneider, M. M., S. A. Buth, B. T. Ho, M. Basler, J. J. Mekalanos, and P. G. Leiman. 2013. “PAAR-repeat proteins sharpen and diversify the type VI secretion system spike.” Nature 500 (7462):350–353. doi: 10.1038/nature12453.

Tang, L., S. Dong, N. Rasheed, H. W. Wu, N. Zhou, H. Li, M. Wang, J. Zheng, J. He, and W. C. H. Chao. 2022. “Vibrio parahaemolyticus prey targeting requires autoproteolysis-triggered dimerization of the type VI secretion system effector RhsP.” Cell Rep 41 (10):111732. doi: 10.1016/j.celrep.2022.111732.

Taylor, N. M., N. S. Prokhorov, R. C. Guerrero-Ferreira, M. M. Shneider, C. Browning, K. N. Goldie, H. Stahlberg, and P. G. Leiman. 2016. “Structure of the T4 baseplate and its function in triggering sheath contraction.” Nature 533 (7603):346–52. doi: 10.1038/nature17971.

Tian, W., C. Chen, X. Lei, J. Zhao, and J. Liang. 2018. “CASTp 3.0: computed atlas of surface topography of proteins.” Nucleic Acids Res 46 (W1):W363–W367. doi: 10.1093/nar/gky473.

Whitney, J. C., D. Quentin, S. Sawai, M. LeRoux, B. N. Harding, H. E. Ledvina, B. Q. Tran, H. Robinson, Y. A. Goo, D. R. Goodlett, S. Raunser, and J. D. Mougous. 2015. “An interbacterial NAD(P)(+) glycohydrolase toxin requires elongation factor Tu for delivery to target cells.” Cell 163 (3):607–19. doi: 10.1016/j.cell.2015.09.027.

Zhang, D., R. F. de Souza, V. Anantharaman, L. M. Iyer, and L. Aravind. 2012. “Polymorphic toxin systems: Comprehensive characterization of trafficking modes, processing, mechanisms of action, immunity and ecology using comparative genomics.” Biol Direct 7:18. doi: 10.1186/1745-6150-7-18.

